# A transcription factor (TF) inference method that broadly measures TF activity and identifies mechanistically distinct TF networks

**DOI:** 10.1101/2024.03.15.585303

**Authors:** Taylor Jones, Rutendo F. Sigauke, Lynn Sanford, Dylan J. Taatjes, Mary A. Allen, Robin D. Dowell

**Author notes:** Contributing authors (Taylor Jones, Rutendo F. Sigauke, Lynn Sanford, Dylan J. Taatjes).

## Abstract

TF profiler is a method of inferring transcription factor regulatory activity, i.e. when a TF is present and actively regulating transcription, directly directly from nascent sequencing assays such as PRO-seq and GRO-seq. Transcription factors orchestrate transcription and play a critical role in cellular maintenance, identity and response to external stimuli. While ChIP assays have measured DNA localization, they fall short of identifying when and where transcription factors are actively regulating transcription. Our method, on the other hand, uses RNA polymerase activity to infer TF activity across hundreds of data sets and transcription factors. Based on these classifications we identify three distinct classes of transcription factors: ubiquitous factors that play roles in cellular homeostasis, driving basal gene programs across tissues and cell types, tissue specific factors that act almost exclusively at enhancers and are themselves regulated at transcription, and stimulus responsive TFs which are regulated post-transcriptionally but act predominantly at enhancers. TF profiler is broadly applicable, providing regulatory insights on any PRO-seq sample for any transcription factor with a known binding motif.

## 1 Introduction

Transcription is a fundamental process that defines cellular function, stress response and cell identity[1]. The regulation of gene expression patterns is driven by a myriad of sequence-specific transcription factors (TFs) that vary in activity based on both cell type and environmental factors. While there are over 1,600 TFs[2] in the human genome, our understanding of how their activity is regulated remains incomplete. For example, there is no consensus on when or where individual TFs are actively altering gene expression patterns.

Transcription factors orchestrate gene regulation programs by altering the activity of cellular RNA polymerases, primarily RNA polymerase II (RNAPII). Some TFs increase transcriptional output (an activator) whereas others decrease transcriptional output (a repressor). Therefore, characterizing when and where TFs are active – not only where they bind in the genome but also when they are actively regulating RNAPII – is necessary to understand their biological function. In fact, one of the goals of the Encyclopedia of DNA Elements (ENCODE) Consortium was to identify all functional regulatory elements in the human genome[3]. In the ENCODE project, the primary method utilized to assess TF activity was chromatin immunopreciptation (ChIP-seq). ChIP informs on the genomic localization of a TF, which reflects the function of its DNA binding domain which typically interacts with DNA in a sequence-specific manner[4].

From ChIP-seq studies, it is possible to infer a position specific scoring matrix (PSSM) for a given DNA binding protein. ChIP-seq studies, however, are low throughput as one sequence-specific protein is evaluated at a time, in one cell type at a time. Furthermore, there is ample evidence that TF binding can occur without altering gene expression[5, 6], as the DNA binding domain is independent of the effector domain (also known as the activation domain or repressor domain). The effector domain interacts with co-regulatory factors that directly or indirectly control RNAPII function to alter gene transcription nearby; thus TFs play crucial role in transcriptional regulation[7, 8].

However, measuring the activity of the effector domain (i.e. measuring TF regulatory activity) has historically been difficult, at least in part because TF regulatory activity can be regulated at multiple stages. For example, TF regulation may occur via changes in protein levels (e.g. TF transcription, translation or degradation) or through post-translational modifications. Many TFs have well-established mechanisms of activation, such as the MAPK pathway phosphorylation events that result in stabilization and activation of MYC [9], or the inhibition of the ubiquitin ligase HDM2 resulting in the stabilization and activation of p53[10–12]. In these cases, the MYC and TP53 genes are present at the mRNA and protein level in most cellular conditions, despite being repressed until activated by specific stimuli. Thus, neither transcription of the gene encoding the TF, nor TF-DNA binding guarantees that it will alter RNAPII transcription. The ultimate outcome of TF effector domain activity is a change in transcription, hence nascent transcription assays are well-suited to inform on effector domain activity.

Run-on RNA sequencing (such as precision run-on sequencing, PRO-seq[13, 14] and global run-on sequencing, GRO-seq[15]) provides a direct read out of RNA polymerase activity as RNA is captured from the actively catalyzing cellular polymerases. These nascent RNA assays have revealed extensive genome-wide transcription, at genes and enhancers[16–19], and demonstrated that most sites of RNAPII initiation give rise to bidirectional transcription. While the function of the resulting RNA transcripts within enhancers is incompletely understood, a technical benefit of these transcripts is that their distinctive profile can be used to annotate active enhancers genome wide[20, 21]. Prior studies on individual TFs found that TF activation resulted in concomitant changes in transcript levels associated with a subset of TF binding sites, as measured by ChIP[22–26]. Subsequent work generalized these findings, showing a strong co-association of TF binding sites with sites of RNAPII initiation, the majority of which occurring at enhancers[20, 21]. The model that emerged was that the regulatory activity of the TF (e.g. activity of the effector domain) results in changes to RNAPII initiation proximal to the TF binding motif[21]. Armed with this result, methods were developed to infer changes in TF activity in response to a perturbation, using nascent transcription data and known TF binding motifs[21, 27–31]. The effectiveness of these methods strongly indicates that nascent transcription serves as a functional readout on the activity of a TF’s effector domain.

Here we sought to develop an algorithm for predicting TF activity from a single nascent transcription experiment. To that end, we develop a statistical framework that compares data from an individual nascent transcription assay to a principled, biologically informed statistical expectation. When a TF recognition motif co-localizes with sites of RNAPII initiation more (or less) than expected by chance, we infer that the TF is functional as an activator (or repressor). Importantly, our algorithm can be used to identify all actively regulating TFs in a single experiment, a technique we call TF profiling. We applied our algorithm to 287 high quality nascent RNA sequencing data sets, representing over 20 different tissues. From this compendium we identify three classes of TFs: ubiquitous (active in all tissue types), tissue-specific, and stimulus responsive. Our method accurately classifies the well known TFs Oct4 and Nanog as active only in embryonic cells. Furthermore, our model uncovered unique sequence features inherent to tissue specific TFs, suggesting a role in the establishment of cell identity.

## 2 Results

### 2.1 An expectation model for TF motif co-occurrences

The activity of the TF effector domain alters nascent transcription proximal to sites of TF binding[7]. Based upon this characteristic, methods to infer TF activity changes from nascent transcription data and TF sequence motifs have been developed[18, 21, 27, 29–31]. One approach derives from a simple metric known as the motif displacement (MD) score, which quantifies co-localization of TF recognition motifs in DNA sequence with sites of RNAPII initiation[21]. The MD-score can effectively identify which TFs are changing in response to specific stimuli. Furthermore, upon stimulation activating TFs showed enrichment of the MD-score whereas repressors show depletion of the score[27], consistent with activators increasing transcription and repressors preventing transcription. These general trends both validate the metric and contribute to its success in differential TF activity inference[21, 27]. Additionally, these results also suggest that MD-scores could be used to infer which TFs were actively participating in regulation within a single experiment.

To test this hypothesis, we sought to develop an algorithm for predicting which TFs were on and actively participating in RNAPII regulation within a single nascent transcription experiment. To that end, we developed a statistical framework for comparing TF motif co-occurrence with sites of RNAPII initiation to a principled biologically informed statistical expectation. When a TF recognition motif co-localizes with sites of RNAPII initiation more (or less) than expected by chance, we infer that the TF is actively participating in RNAPII regulation as an activator (or repressor). Consequently, we call our algorithm TF profiling.

Consider the MD-score metric in a rigorous mathematical framework. Let *X_k_*= *µ*_1_,*µ*_2_,…, *µ_n_*be the RNAPII initiation sites (*µ*) for a set of bidirectional locations genome-wide for some experiment *k*. Importantly, sites of bidirectional transcription can be identified directly from nascent transcription data[21, 29, 32, 33]. Let *Y_j_* = *y*_1_,*y*_2_,…, *y_m_* be the set of all significant motif instances for some TF-DNA binding motif model *j* genome-wide, which is invariant given the genome of interest (Figure 1A). We can then plot the motif displacement distribution (Figure 1B) as a heatmap, where heat indicates the number of motif hits (*Y_j_*) relative to the sites of RNAPII initiation (*X_k_*). In this framework, we can calculate the MD-score as:

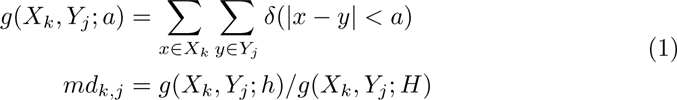

where *g*() quantifies the count of motif hits for a given motif (*j*) across the complete set of RNAPII initiation sites (*X_k_*). The *δ*(.) term is a simple indicator function that returns one if the distance between one RNAPII initiation position (*x*) has an instance of the TF-DNA binding motif (*y*) within a specified distance (*a*). Hence, the MD-score (*md_k.j_*) for a given experiment *k* and TF recognition motif *j* quantifies co-localization of motif instances near sites of RNAPII initiation (*h* = 150bps) relative to a larger local window (*H* = 1500bp). Importantly, our prior work showed that the value of the MD metric depends on precisely defining sites of RNAPII initiation, which is readily accomplished in nascent transcription assays[27]. In fact, for a given TF, the precision of motif co-localization with RNAPII initiation is comparable to TF motif localization with ChIP-seq. We emphasize that although H3K27ac ChIP-seq data could be considered a proxy for genome-wide RNAPII initiation regions at enhancers[27], the regions are too broad to get reliable information about TF binding sites. Therefore, nascent transcription (e.g. PRO-seq) strikes the balance for defining TF activity with precision (comparable to TF ChIP) and scale (comparable to H3K27ac ChIP; Supplemental Figure 1A-C).

**Fig. 1.**
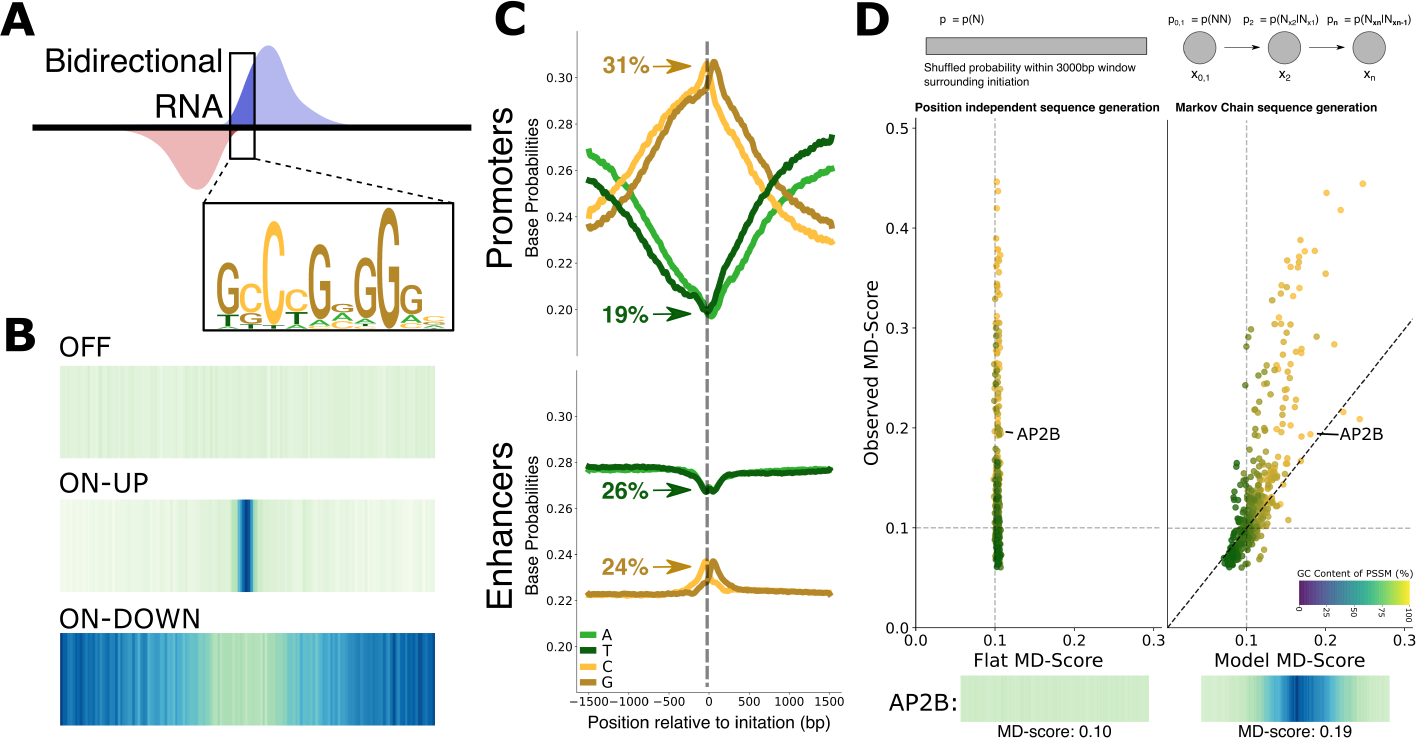
Overview of the TF profiling model. A) Cartoon representing the co-localization between bidirectional transcription observed in nascent RNA sequencing (blue and red are data on each strand) and TF motifs. The PSSM for AP2B is shown. This co-localization can be used to assess global motif displacement scores. B) Heat maps representing the motif displacement distribution[21] for three distinct TFs with different activation states, OFF (ZN586), ON-UP (SP3) and ON-DOWN (PAX5). The center of the heat map is the position of the middle of the bidirectional (PolII initiation site) and the heat (darker is more) represents the number of motif instances at that position (relative to the center) genome-wide. C) Observed promoter (top) and non-promoter(e.g. enhancers, bottom) per position base probabilities surrounding PolII initiation sites show a profound GC bias. In this data set[75], bidirectionals are 30% at promoters (top) and 70% at non-promoters (enhancers, bottom). D) The observed motif displacement score distribution assuming a flat background and no positional information (left) compared to a position dependent di-nucleotide Markov background (right). Each dot is a single TF position specific scoring matrix, colored by its inherent GC content. The probability (*p_i_*) is defined by the observed probabilities (N) at position *i*. The position and motif displacement distribution for AP2B is shown with both background models.

To utilize the MD-score as a metric for TF activity, we seek to calculate an odds ratio: the MD-score observed (in a single experiment) compared to the expected MD-score (from a statistical model). Conceptually, the expected MD-score must reflect the nucleotide biases of not only the TF-DNA binding motif but also the distinct non-stationary patterns of sequence inherent in genomes. In particular, mammalian genomes have GC-content enrichment at promoters[3, 34] and enhancers[21], consistent with sites of RNAPII initiation. Gene promoters are associated with open chromatin and are highly enriched for CpG islands[35–37]. Whereas the human genome is approximately 60% AT, promoters are approximately 60% GC and enhancers are more modestly GC rich, reaching a nearly equal composition of all four bases (Figure 1C; see Methods for promoter and enhancer classification). The difference between enhancer and promoter GC content is statistically significant (Supplemental Figure 2). Importantly, in both cases (enhancers or promoters) the bias is position dependent, reaching a maximum bias coincident with the inferred position of RNAPII loading (*µ* in our model). We observe that this bias correlates with the overall transcription level, where regions with higher transcription levels tend to display a higher GC content over a broader initiation region (Supplemental Figure 3). Because of the positional base composition bias at RNAPII initiation regions, certain motif instances will be favored (high GC) or disfavored (high AT) by chance alone. Our background expectation model must account for this inherent bias.

Therefore, we took a simulation based approach to the development of the expected MD-score. Specifically, we leverage a dinucleotide model of positional nucleotide preference (Figure 1D), which accounts for known genomic dinucleotide biases, such as the general preference for CG in CpG islands compared to GC (Supplemental Figure 4). To this end, sequences of the length of 2*H* nucleotides were generated, accounting for dinucleotide preferences in regions of RNAPII initiation. Importantly, the positions *i* are defined relative to the RNAPII initiation position *µ* (e.g. the generated sequence is *µ±* H). Let *x_n_*= *x*_1_,*x*_2_,…*x*_2_*_H_* where the probability of a specific nucleotide at each *x_i_* is determined based on the nucleotide *x_i−_*_1_. Thus each position is described by the conditional probability *p*(*N_x__i_ |N_x__i−_*_1_), where *N* represents one of the four nucleotides (A, T, C or G). The initial dinucleotide *x*_1_*x*_2_ is calculated as *p*(*N*_1_*, N*_2_) and all sub-sequent positions are based on the conditional probability of the previous position. Therefore, we generate the sequences as:

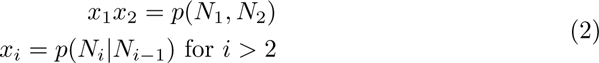

Importantly, we further capture the natural diversity in GC bias (both magnitude and width) by simulating from distinct promoter and enhancer dinucleotide probabilities (Supplemental Figure 4). The proportion of bidirectional calls at promoters (versus enhancers) varies across data sets (Supplemental Figure 5), which may be biological or could reflect ascertainment biases since promoters tend to be more highly transcribed. Since promoters are considerably more GC rich than enhancers (Supplemental Figure 2), TFs with GC rich motifs will be disproportionately enriched (false positives) in data sets with high promoter content. To control for this, we simulated sequences from the two classes (enhancers and promoters) in proportion to the observed ratio for a total of 10^6^ instances (see Methods). Using these simulated sequences we calculated expected MD-scores (Equation 1). This enables us to compare the expected (i.e. model derived, x-axis) to observed (i.e. experimentally observed, y-axis) MD-score for a single data set[38] as shown in Figure 2A. Thus, the expectation model is calculated on a per data set basis to accurately reflect the composition of initiation regions inherent to that cell type and condition.

**Fig. 2.**
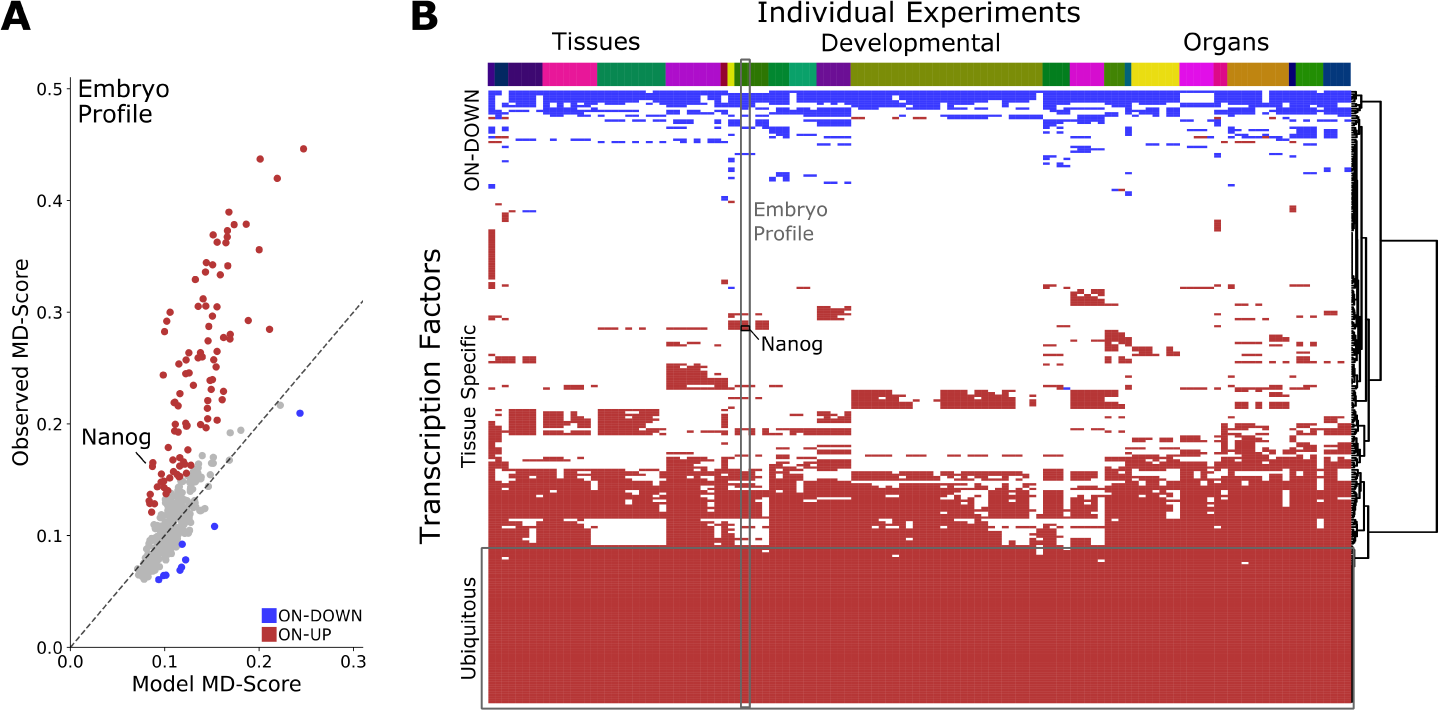
Generating and clustering of TF profiles. A) Scatter plot showing the expected (x-axis) and observed (y-axis) MD-scores for all PSSMs in HOCOMOCO for for embryonic stem cells[75]. Significant differences are colored with ON-UP red and ON-DOWN blue (see Methods). The collection of ON and OFF (grey) labels is the TF profile for this experiment. B) Ward clustering of TF profiles (columns) in 126 samples representing over twenty tissue types. Each sample is labeled by its tissue of origin (top, colored bar) with tissue labels further classified into tissues (e.g. bone, blood), developmental (e.g. fetal, embryo) and organ (e.g. breast, kidney). The ON-DOWN TFs (at the top, blue) tend to be shared across samples. The ON-UP (red) shows a variety of patterns including tissue specific pockets (middle) and ubiquitously on and active (bottom).

### 2.2 Building TF activity profiles across tissues

The next step was to assess the statistical significance of TF activity for each TF-PSSM occurrence; that is, to statistically ask which motif hits in Figure 2A were significantly more (or less) co-localized with RNAPII initiation sites than our background model suggests (versus expectation). Logically, TF-motifs with greater (or less) than expected co-localization are the TFs we infer as ON (ON-UP and ON-DOWN, respectively) and participating actively in RNAPII regulation.

To this end, the MD-scores for 388 TF motifs (HOCOMOCO core version 11[39]) were calculated for all control data sets of sufficient quality (n=126; see Methods). In each data set, the observed MD-score was compared to the expected MD-score. To assess statistical significance, we further assumed that the majority of TFs will be OFF across all control data sets (75% not significantly different from expectation; Supplemental Figure 6A). The distribution of residuals (Supplemental Figure 6B) was then used to assess significance for all TF motifs within all control data sets. This resulted in a range of 80-164 TFs that were called ON in any given data set (mean=123.5, p-value *<* 0.05). The ON TFs can be split into two categories, on and enriched (ON-UP, activators; range of 74-148, mean=109.9) or on and depleted (ON-DOWN, repressors; range 5-28, mean=13.6) (Supplemental Figure 6C). We refer to the collection of ON TFs (either UP or DOWN) for a given cell line as its TF activity profile.

An example TF profile is shown in Figure 2A, where red and blue represent TFs that are classified as ON-UP (red) and ON-DOWN (blue) and grey represents TFs that are OFF. When applied to an embryonic stem cell data set (Figure 2A; n=3 biological replicates)[38], we called 95 enriched and 9 depleted TFs (Fig. 2A). Enriched TFs included the pluripotent factors responsible for embryonic stem cell self-renewal, Oct4 (pval=1.3*e^−^*^5^), Nanog (pval=2.7*e^−^*^5^) and SRY-Box Transcription Factors 3 and 4 (pval=0.008, pval=0.03 respectively). Importantly, across 3 additional independent embryonic stem cell data sets[40–42] the same pluripotency factors were consistently called as active. TFs identified as depleted include a variety of known repressors including SNAI2, CEBPA and E2F1[43–45].

We next sought to expand our examination of TF activity profiles by clustering the profiles across tissues and cell types that had high quality nascent RNA sequencing data (See Methods). In total, we examined 126 distinct data sets representing a total of 299 nascent RNA-sequencing samples in basal conditions (i.e. normal growth or control samples). We used Ward’s method to cluster the TF activity profiles (using Euclidean distance) across the DBNascent high-quality control samples. We found that the major determinant in clustering was tissue identity (Figure 2B; Supplemental Figure 7).

Moreover, the clustering of TF profiles suggested that at the extremes, some TFs are active across nearly all cell types and other TFs are tissue specific. Notably, only 4-6 TFs per tissue type were truly tissue specific, however, they were the major determinant for clustering. For example, the TF MyoD is a strong determinate in muscle differentiation[46] and was uniquely ON-UP (pval= 7.0*e^−^*^9^) in the myoblast data set[47] and OFF in the other 125 data sets tested. The blood associated factor, GATA-2, was uniquely called as ON-UP across several blood samples[29, 48–52], and was notably OFF within the other data sets. In addition to these well known cell type specific TFs, we also recovered less well annotated TFs that infer uncharacterized biological functions. For example, ZNF121 in blood, or ZNF146 in organ function.

We also noticed that some TFs implicated in general cellular processes were commonly called ON-UP in the TF profiles. These “ubiquitously active” TFs included members of the ETS family, the E2F family, and KLF/SP family. These TFs have highly redundant binding motifs and their biological functions relate to cellular homeostasis and proliferation[53–55]. Additionally, these ubiquitous TFs may help maintain promoter accessibility and/or enable promoter-promoter looping[56, 57].

### 2.3 TF region selection across tissues

We next sought to further characterize the two extreme classes of TF regulatory activity: the ubiquitous and tissue-specific classes of TFs. First, we noted that the GC content of the TF recognition motifs differed between ubiquitous and tissue-specific TFs (Figure 3A). The ubiquitous TFs bound GC-rich regions that were close to the average GC composition at promoters, consistent with prior reports[56]. By contrast, the tissue-specific TFs tended to have motif preferences closer to genomic background (Figure 3A). Notably, this result was recapitulated in SELEX and protein binding microarray data independent of genomic context (Supplemental Figure 8).

**Fig. 3.**
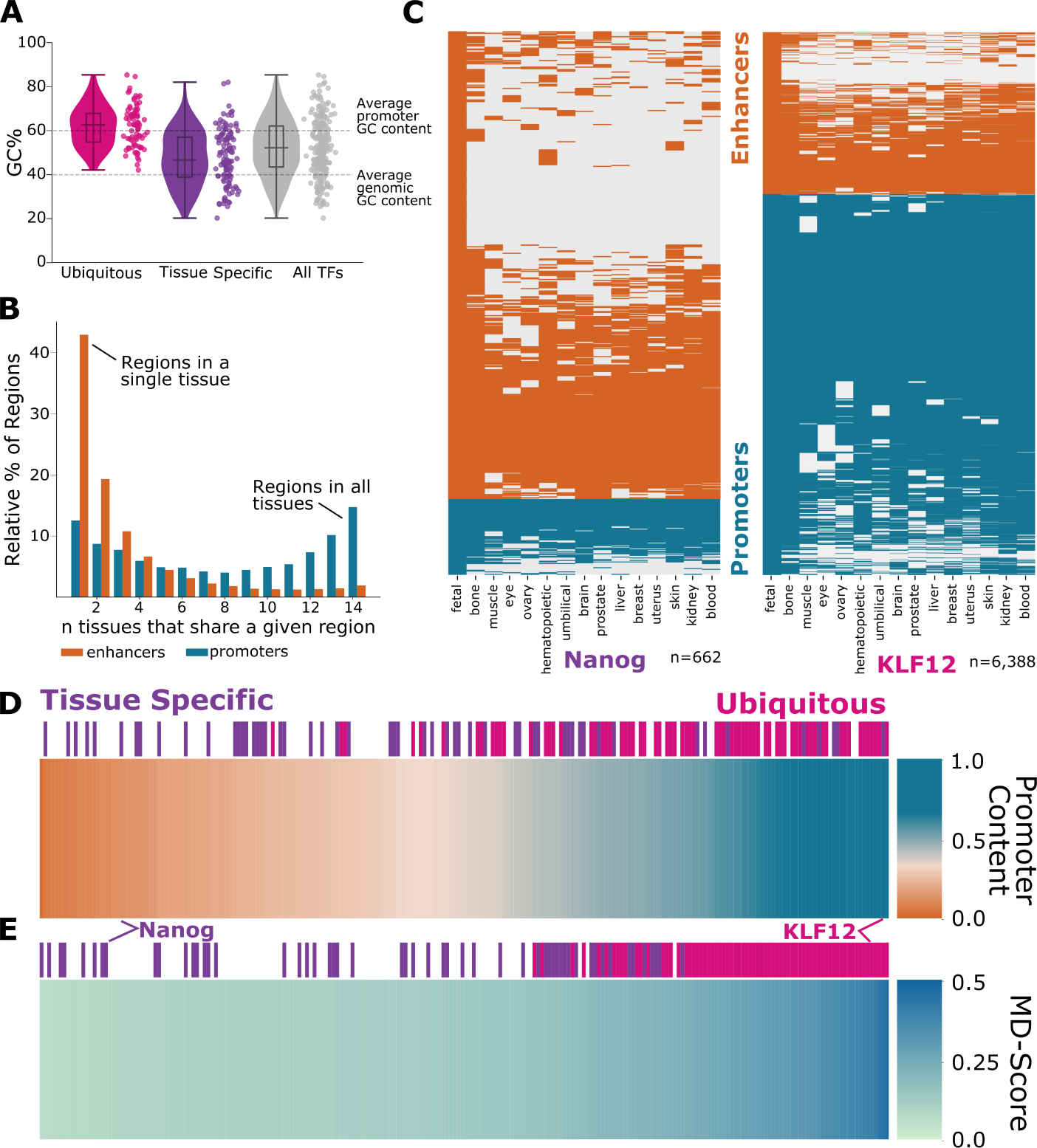
Tissue specific and ubiquitous factors have distinct localization and regulation preferences. A) Violin plots showing the GC content of the TF motif (PSSM) of ubiquitously shared TFs (pink), tissue specific TFs (purple) and all TFs (grey). B) Bar plot showing that enhancers (orange) are far more tissue specific than promoters (blue) which tend to be on in all tissues. C) Set of consensus bidirectional regions in embryonic stem cells containing a centered motif for the tissue specific Nanog (left) or the ubiqitous KLF12 (right), colored (promoter: blue, enhancer: orange) by presence or absence across multiple tissues (x-axis). Across 662 regions containing Nanog in ESC cells, 86.1% of regions are enhancers, with some being shared (bottom, solid orange) and others being more tissue specific (middle, mostly white). Across the 6,388 KLF12 containing regions in ESCs, 70.3% are promoter regions. D) Fraction of ChIP-seq binding sites at the 5*^′^* end of genes (promoter, blue) or at distal regulatory regions (enhancers, orange) for ubiquitous TFs (pink) and tissue specific TFs (purple)[61, 62]. E) Motif displacement score as a heat map (darker is higher) for ubiquitous TFs (pink) and tissue specific TFs (purple).

Given the sequence preferences inherent to the ubiquitous and tissue-specific TF classes, we next wondered whether these TFs would act at distinct genomic regions (e.g. promoters vs. enhancers). As previously noted[58–60], enhancer regions are more tissue specific whereas promoters are often transcribed more broadly across cell types (Figure 3B). Thus, we examined the number of enhancer and promoter regions contributing to each TF’s activity profile. We observed that tissue specific TFs predominantly regulate at enhancers whereas ubiquitous TFs generally regulate at promoters (Figure 3C). For example, Nanog is an embryonic specific TF. The vast majority of transcribed regions containing the Nanog motif in embryonic stem cells (86.1%) are enhancers, the majority of which are unique to the embryonic tissue samples. In contrast, the regions with the KLF12 motif, a ubiquitous TF, tend to be promoter associated (70.3%) and transcribed across most tissues.

We next wondered whether the region bias identified by TF profiling was also present in transcription factor chromatin immunoprecipitation (ChIP). Thus we next examined TF ChIP-seq data curated from cistromeDB[61, 62]. To select for high quality ChIP signal, we only considered TF ChIP-seq peaks within regulatory regions (n=53,244 promoters, n=559,150 enhancers). We then asked how often ChIP peaks fell within promoter regions versus enhancer regions. The ChIP data further supported the observation that ubiquitous TFs bind and regulate predominantly at promoters, whereas tissue-specific TFs bind and regulated predominantly at enhancers (Figure 3D). This result is reliably captured by the MD-score approach, where ubiquitous TFs have higher MD-scores on average than tissue specific factors (Figure 3E).

### 2.4 Regulation at Transcription

Given the distinct binding sites and biological functions of the ubiquitous and tissue-specific TFs classes, we next asked whether the regulation of these TF classes was distinct. To this end, we first examined the transcription level of the gene encoding each TF. For example, we assessed the transcription level of GRHL2 (tissue specific) and CREB1 (ubiquitous) across the control data sets. The tissue specific TF had many samples with low gene transcription and a few samples with high gene transcription. Thus, the distribution of the TF gene transcription level followed an exponential distribution, consistent with an transcription of the gene in a limited subset of the data. In contrast, the transcription of the ubiquitous TF was normally distributed (Figure 4A), consistent with the TF gene being transcribed in all samples. Consequently, we classified each gene encoding a TF as either fitting an exponential or normal distribution. Notably, the tissue specific TFs tended to fit an exponential distribution, like GRHL2, and ubiquitous TFs had a bias towards a normal distribution of transcription like CREB1 (Supplemental Figure 9). In sum, the two classes show distinct corresponding cumulative density functions for TF transcription across a subset of high confidence TFs (Figure 4B; see Methods).

**Fig. 4.**
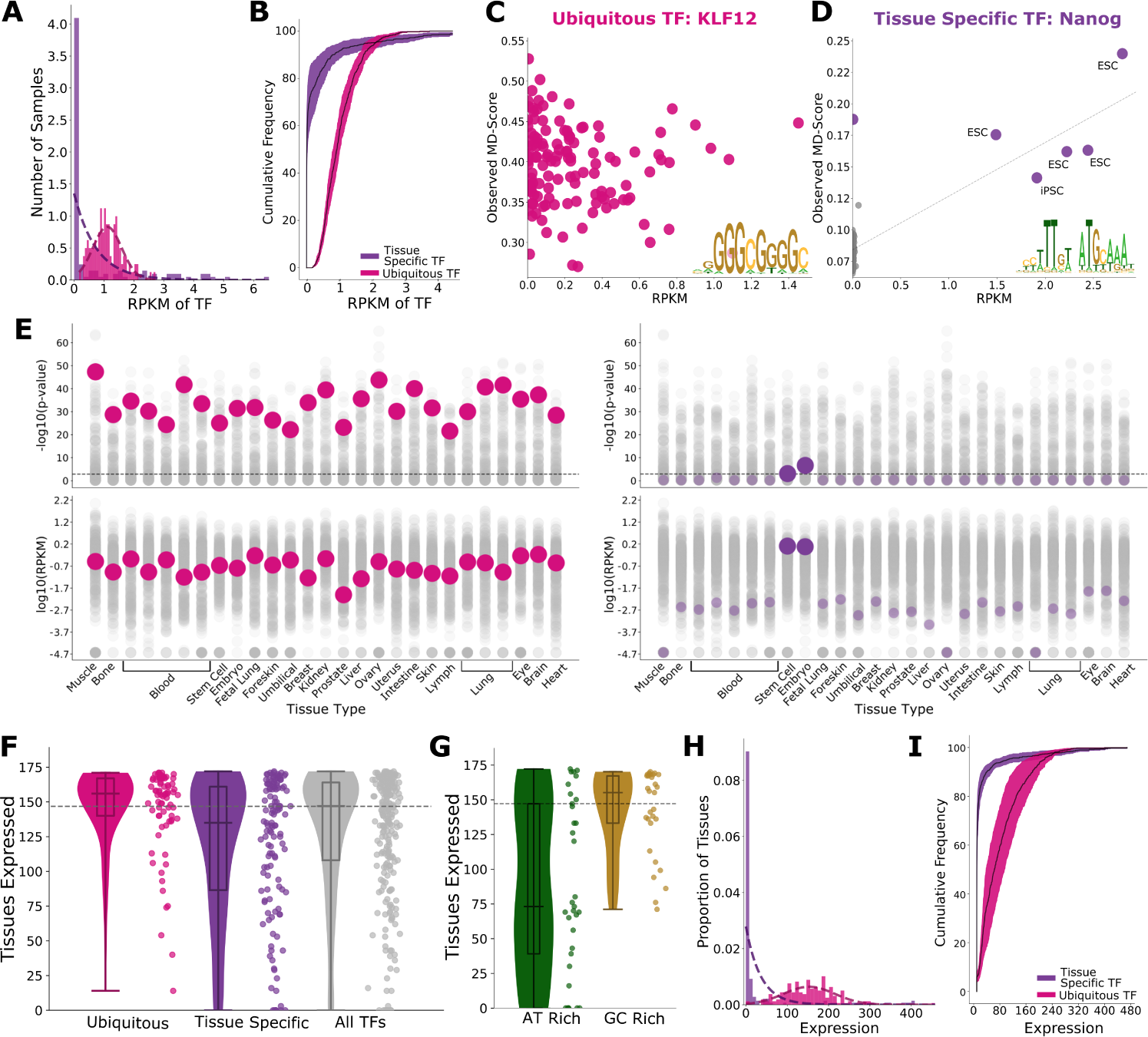
Tissue specific TFs are regulated at transcription. Ubiquitous factors are post-transcripionally regulated. A) Histogram of the transcription level of a tissue specific TF (GRHL2; purple) and ubiquitous TF (CREB1; pink) in nascent RNA-seq. B) Cumulative distribution function of the transcription of the gene encoding the TF (RPKM) for a set of high confidence tissue specific factors (purple) and ubiquitous factors (pink) across 126 control experiments. The relationship between the transcription level of the gene encoding the TF (x-axis) and observed MD-score (y-axis) for C) ubiquitous TF (KLF12) and D) a tissue specific TF (Nanog). HOCOMOCO PSSMs shown in lower right corner. E) Plot of the significance of the MD-score (top) and the transcription of the gene encoding the TF (bottom) for all TFs (grey), highlighting KLF12 (left, pink) and Nanog (right, purple). F) Violin plots of frequency of expression in single cell RNA-seq[63] across 172 tissues for ubiquitous (pink), tissue specific (purple) and all TFs (grey). G) Violin plots of frequency of expression in single cell RNA-seq[63] across 172 tissues for TFs with the bottom 10% (green) and top 10% (gold) GC content within their PSSMs. H) Histogram of the number of tissues that a tissue specific TF (GRHL2; purple) and ubiquitous TF (CREB1; pink) are expressed in by single cell RNA-seq[63]. I) Cumulative distribution function of the steady-state RNA level (scRNA-seq) for the same high confidence tissue specific factors (purple), and ubiquitous factors (pink) across 172 tissues from atlas of fetal gene expression[63].

We next examined these distributions the context of TF regulatory activity (the MD-score). As a representative example, Figure 4C shows a plot of the MD-Score vs. the transcription level (RPKM) for the ubiquitous TF KLF12. There was no correlation between the TF gene transcription level and predicted TF activity. This pattern was observed across many ubiquitously shared TFs such as SP1 and ETV1 (Supplementary Figure 10A) and suggests that there is no obvious relationship between ubiquitous TF transcription level and its activity (i.e. MD-score). In contrast, the tissue specific TF Nanog shows a positive correlation between its activity (MD-Score) and gene transcription level (RPKM; Figure 4D). Moreover, this positive correlation was observed for many tissue specific factors, including MyoD and GATA-2 (Supplementary Figure 10B). This result indicates that tissue specific TFs are not transcribed unless they are actively regulating within a cellular context, suggesting that repression of tissue specific TFs transcription plays a role in blocking their function. In summary, the two classes of TFs, ubiquitous and tissue specific, have categorically distinct transcription patterns, and suggests biologically distinct mechanisms of TF gene activation (Figure 4E).

To further probe these results, we sought to determine whether these trends would be recapitulated from steady-state RNA levels. We utilized single cell RNA-seq (scRNA-seq) data from atlas of fetal gene expression[63] as it allowed us to capture expression values for these TFs across 172 distinct human tissues. We observed that the ubiquitous TFs were generally expressed in all tissue types whereas the tissue specific TFs were expressed in fewer tissues (Figure 4F). When we assessed the expression of TFs with the lowest GC content motifs (bottom 10%) we found that the median number of tissues with TF expression falls far below the total median. 72.7% of TFs with the low GC motif set are classified as tissue specific. The TFs with the highest GC content motifs (top 10%) are expressed in more tissues than expectation and many are ubiquitously active TFs (65.5% ubiquitous; Figure 4G). This was also reflected by their classified distributions of expression (exponential or normal). Similar to the observed trends in nascent transcription, tissue specific TFs are expressed in fewer tissues lending to an exponential fit, whereas the ubiquitous TFs have gene expression that tends to be normally distributed (Figure 4H-I). Overall, the trends we observed at the transcriptional level (PRO-seq) are recapitulated at the steady-state RNA level (scRNA-seq) suggesting this is a fundamental regulatory strategy for ubiquitous TFs versus tissue specific TFs.

### 2.5 Stimulus Responsive TFs

The ubiquitous and tissue specific TFs represent the extremes of ON and OFF patterns within our clustering (Figure 2B. Yet many transcription factors were ON in groups of samples, either several tissues or more sporadically across samples. We reasoned that stimulus responsive TFs could give rise to a more sporatic pattern of activity, as the activity of the TF would depend on the fine details of the growth environment. Thus we next sought to identify high confidence stimulus responsive TFs. To accomplish this, we identified 161 data sets in treatment conditions from corresponding publications with our control data sets[60]. We applied TF profiling to this “perturbation” collection (Supplemental Figure 11A-C), identifying 53 high confidence stimulus responsive TFs. We next sought to characterize the 53 high confidence stimulus responsive TFs.

We first probed whether the stimulus responsive TFs have a have recognition motif preferences comparable to either the ubiquitous or tissue specific TFs. We determined that stimulus responsive TFs have recognition motifs that are similar to genomic background, as seen with tissue specific TFs (Supplemental Figure 13A). We next examined ChIP-seq data for the stimulus responsive TFs, finding that they bind and act primarily in enhancer regions, similar to tissue specific factors (Supplemental Figure 13B-C). Among each of our classified TF groups, we found a positive correlation between the GC content of the recognition motif and the preference for binding within promoter regions, where ubiquitous TFs dominate the high GC percentage regime and the other two classes (tissue specific and stimulus responsive) behave similarly in the low GC percentage regime (Figure 5A).

**Fig. 5.**
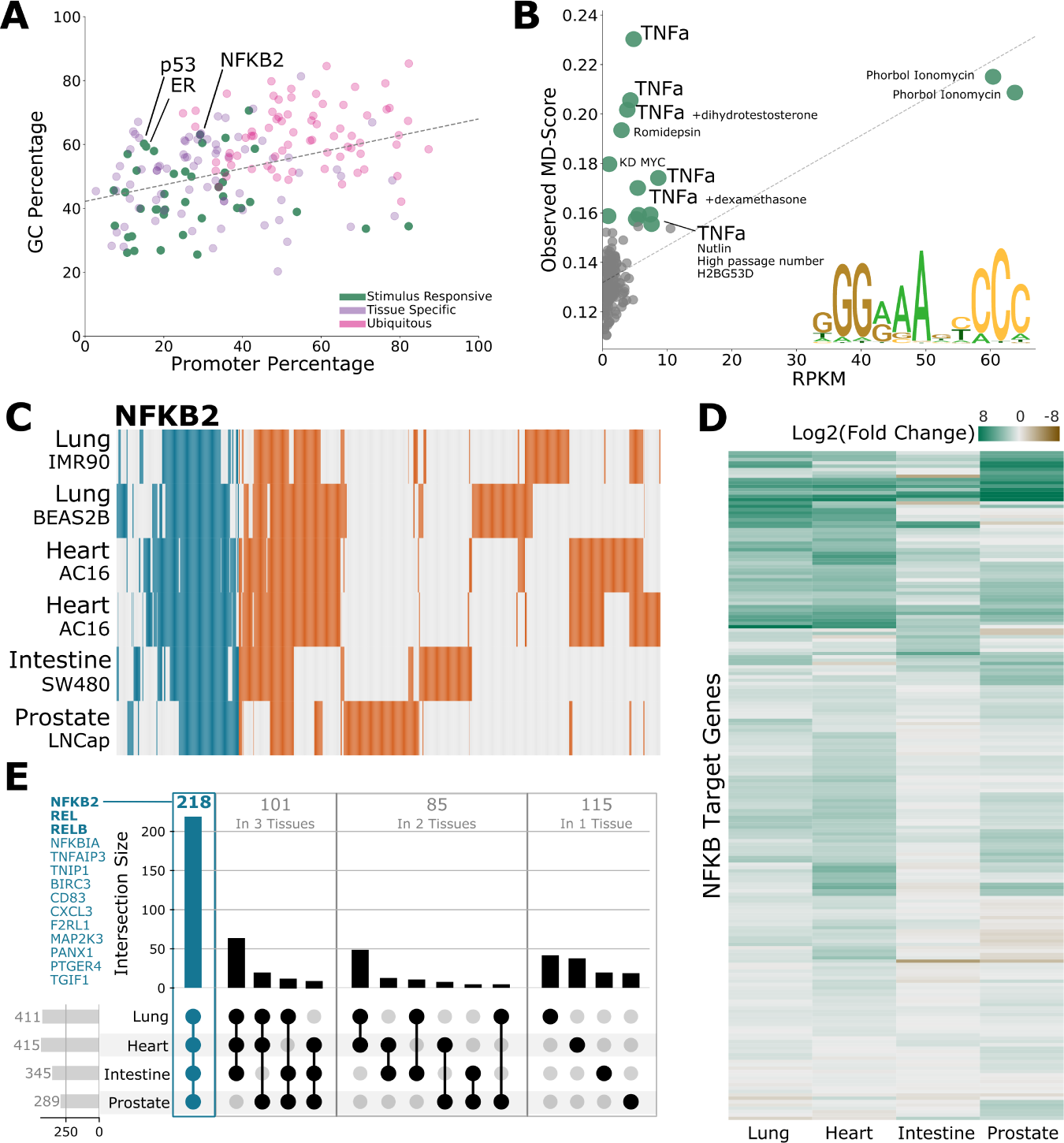
Stimulus responsive TFs utilize distinct regions to achieve equitable stimulus response across tissues. A) Scatter plot of the percentage of ChIP-seq sites within promoters (x-axis) vs the GC content of the TF motif (PSSM, y-axis) for ubiquitous (pink), tissue specific (purple) and stimulus reponsive (green) TFs. B) Comparing the transcription level of the gene encoding the TF (x-axis) and MD-score (y-axis) for the stimulus responsive TF NF*κ*B. Significant ON-UP instances of NF*κ*B are colored green and labeled with the stimulus. Inset it the PSSM for NF*κ*B2. C) Bidirectional regions with a centered NF*κ*B2 motif from TNF*α* treated cells (larger font in panel B, four tissues). Color indicates detection of the bidirectional, with enhancer regions in orange and promoters in blue. Across 2,302 regions with NF*κ*B2 motif instances, 77.5% of these regions are classified as enhancers. Of the enhancers, 1,406 (78.9%) are unique to a given tissue type. Other subunits of NF*κ*B shown in Supplemental Figure 14. D) Upset plot of promoter regions (n=519) shown in panel C where 42.0% are shared across all tissues and 81.0% are shared between at least two tissues. In the set of promoters shared across all tissues (teal) there are numerous NF*κ*B target genes (teal text). Three target genes (NF*κ*B2, REL and RELB, bold) are three subunits of the NF*κ*B TF complex. E) Heat map of NF*κ*B target genes (y-axis) across the four tissues in panel C(heart, intestine, lung, prostate). Up-regulated (green); gold (down-regulated).

We next examined the regulation of the gene encoding the stimulus responsive TFs (Figure 4C,D, Supplemental Figure 10A,B). Intriguingly, we found that many stimulus responsive TFs are broadly transcribed, similar to ubiquitous factors, but active in only a subset of samples, similar to tissue specific TFs (Figure 5B, Supplemental Figure 12A-C). This is consistent with the fact that many stimulus responsive factors are post-transcriptionally regulated. For example, under normal conditions, p53 is constantly transcribed and translated, but subsequently degraded via the ubiquitin ligase HDM2[64, 65]. Consistent with post-translational regulation, we observed elevated activity scores only in samples where p53 was directly stimulated (Supplemental Figure 12B).

To fully understand the distinct behavior of stimulus responsive TFs, we selected NF*κ*B as a case study, as it has the most high-quality data (six data sets, four tissue types[26, 66–70]). NF*κ*B is a key regulator of the inflammation response across tissue types[71]. Across four of the NF*κ*B subunits (REL, RELB, NF*κ*B2 and T65) we noted that the TF region selection differed across tissue types, specifically within enhancer regions which represent the majority of putative binding sites (Figure 5C, Supplemental Figure 14A-C). In fact, within a tissue with multiple cell lines (Lung; IMR90 and BEAS2B) the region selection varied, but within a tissue with the same cell line replicated from different publications (Heart; AC16) the enhancer region selection was consistent. This suggests that the NF*κ*B response regions are defined by the cell type. Despite this, there was a robust NF*κ*B response in all tissues (Figure 5D, Supplemental Figure 15). While enhancer region selection was highly variable, active promoter regions with NF*κ*B2 motifs are more consistent across tissues. Out of 519 promoter regions across the TNF*α* treated samples, 218 (42.0%) are shared across all tissues (Figure 5E). We next examined the genes associated with these promoter regions. Multiple genes were direct NF*κ*B targets, including subunits of the NF*κ*B TF (NF*κ*B2, REL and RELB).

## 3 Discussion

Here we present TF profiling, a method of TF activity inference that identifies which TFs, among hundreds with well-characterized motifs, are actively regulating RNAPII from a single nascent RNA-sequencing experiment. The method relies on a robust sequence based expectation model derived from the base probabilities at RNAPII initiation regions. Using this method, we can identify which TFs are ON and active, regardless of whether the TF is an activator or repressor. We anticipate that this method will be broadly useful for assessing the set of TFs active in any cell type, provided that high-quality nascent sequencing data is available. Interestingly, the TF profiling method identified 3 classes of TFs: ubiquitous TFs which are always on regardless of cell type or condition, tissue specific TFs that drive cell identity and stimulus responsive TFs that are poised to alter transcription in response to a perturbation. We also showed that these TF classes have distinct DNA binding preferences and are regulated via distinct mechanisms. Because TFs drive all biological processes and are among the most important class of proteins in biology, it is critical to develop tools to reliably assess TF activity.

The ubiquitous TFs have GC-rich recognition motifs and bind preferentially at promoters. The ubiquitous TFs are represented in part by the ETS, KLF, E2F, ATF, and SP1 families. We note that among the ubiquitous TFs (n=78), many motif preferences are similar and therefore difficult to distinguish from each other. While there may be subtle differences in which TFs are active in a given cell line, a subset of these ubiquitous TFs are always active regardless of cell line or condition. In agreement, most of the ubiquitous TFs were transcribed in nearly all data sets tested, suggesting they function cooperatively or redundantly. Moreover, individual ubiquitous TFs are typically not essential, suggesting they behave collectively to regulate RNAPII function, perhaps to help maintain nucleosome-free promoters[56], though genomic regions with high GC content naturally exclude nucleosomes[34]. Finally, ubiquitous TFs regulate genes important for cellular proliferation, metabolism and homeostasis[72–74], consistent with their general requirement across cell types.

Distinct from the ubiquitous TFs, the tissue specific TFs preferentially bind enhancers with binding motifs that have a nucleotide composition similar to genomic background, i.e. more AT-rich. Importantly, there are a subset of TFs that bind AT-rich regions but were not called ON in any of our data sets. Many of these never ON TFs are implicated in cell identity for cell types with no nascent RNA sequencing data. For example, UNCX is a TF implicated in regulation of the cerebellum with an AT-rich binding preference. Yet no cerebellum data is present within DBNascent, which could explain why we do not see UNCX as ON in any of these data sets. Many of the tissue specific group of TFs are not transcribed unless they are ON within a given cellular context, e.g. their activity may be regulated by their transcription.

Many TFs are neither ubiquitous or tissue specific. This includes TFs that are on in subsets of, often related, tissues. It also includes the stimulus responsive TFs, which share many of the same recognition properties as the tissue specific TFs. Namely, they bind predominantly at enhancer regions and have recognition motifs similar to genomic background composition. Yet unlike tissue specific TFs, the stimulus responsive TFs are typically transcribed across a broad range of tissue types and conditions (similar to the ubiquitous TFs). This pattern is consistent with post-transcriptional regulation of these TFs, allowing them to be poised for activation but are not always ON; instead, post-transcriptional mechanisms regulate their activity.

In sum, we discovered 3 distinct classes of TFs: ubiquitous, tissue specific, and stimulus responsive. We speculate that the binding preferences and mechanism of regulation for a given TF may be predicted based on the TFs function. While it’s known that tissue specific TFs play a crucial role in defining cell identity, we postulate that these TFs are not transcribed unless actively regulating transcription as they are key players in establishing tissue specific enhancer regions. Furthermore, it is tempting to speculate that these tissue specific enhancer patterns would then directly explain the subset of tissue specific stimulus responsive regulatory sites. However, the set of tissue specific TFs are not enriched directly in or adjacent to the cell type specific stimulus responsive sites. This contradiction suggests that tissue specific stimulus responses may arise from some complex interaction between tissue specific TFs at some sites and other factors such as chromatin state or transcriptionally active domains.

## 4 Methods

### Code availability

The stand-alone TF Profiler application can be found on github (https://github.com/ Dowell-Lab/TF profiler). TF Profiler takes an annotation file for bidirectional regions from a nascent sequencing experiment and derives a TF profile. This includes generating simulated sequences based on the base composition of the regions provided, scanning for PSSM hits within the genome and statistically assessing TF enrichment and depletion.

Additional stand-alone scripts and useful data files associated with this work can be found on github (https://github.com/Dowell-Lab/TF profileradditonal scripts).

### Curating data from DBNascent

All nascent RNA sequencing data, which includes both precision run-on sequencing (PRO-seq) and global run-on sequencing (GRO-seq), were obtained from DBNascent[60]. Briefly, the database contains 502 human PRO-seq samples from 60 publications and 780 human GRO-seq samples from 106 publications. All data were aligned to the human reference genome (hg38), subjected to extensive quality control, and processed for identifying sites of bidirectional transcription (also known as transcribed regulatory elements) using standardized Nextflow pipelines, as described in Sigauke et al.[60].

To prepare the data for TF profiling, high quality nascent RNA samples were selected from the database, with minimum quality score of 4 (minimum of 5 million reads, over 50% of reads map to the reference genome and less than 95% duplication). Within these samples, Tfit[17] was utilized to annotate bidirectional regions. All Tfit calls between biological replicates of a given cell type within a given paper were merged using muMerge[27]. If only one biological replicate passed the quality score cut-off it was still used for subsequent analysis as a single replicate data set. The grouped biological replicates within a cell type and paper are referred to as “data sets” in this study. Bidirectional annotations were merged on a per-cell and per-paper basis to maximize number of reliable calls per data set.

To account for low complexity in some data sets, which can arise from either poor pull down efficiency or high sequencing noise, we also filtered data sets based on the quality of the bidirectional calls. To this end, we required that the region within 2*h* of *µ* had a base composition of at least 50% GC content. The final requirement is that at least 50% of called regions must not be at an annotated promoters. If promoter regions are over-represented then we lose sensitivity when calling many TFs, as most TFs bind predominantly at distal regulatory elements, such as enhancers.

### Curating a master bidirectional region list

After this two step quality control process, we ended up with 126 distinct data sets from 88 publications that represent 79 unique cell lines under basal conditions (e.g. basal, normal growth conditions; n=299 unique biological samples). Samples were merged step-wise, with all samples of a given tissue type were merged into a tissue specific regions file. Any region less than 20nt were windowed to be at least 20nt in length. The tissue specific region files were then merged into the master file. From the same publications as the control samples an additional 161 data sets with identifiable perturbation or genome modified conditions also passed the quality control process. These samples come from 65 of the 88 control publications and represent 46 unique cell lines (n=411 unique biological samples).

Regions within the master file (control samples only) were divided into two sets: promoters and enhancers. Promoters were defined as all regions (windowed by *h*=150) within 1000bp (300bp upstream, 700bp downstream) of RefSeq (hg38 release GCF (109.20190607 2019 06) annotated transcription start site (TSS). All other regions were labeled as enhancer. This resulted in a total of 53,244 promoter regions, and 611,963 enhancer regions within the master file.

### Calculating positional probabilities

Both regions types (promoters and enhancers) used to independently extract two sets of positional probabilities surrounding *µ* using a window size of *H* = 1500 (e.g. *µ ±* 1500). Sequences were extracted using bedtools getfasta (bedtools/2.25.0). All ambiguous bases were replaced with randomly sampled nucleotides (A, C, G, T) using a flat distribution (all bases equal probable). Two distinct probability distributions are then tallied from the sequences. First, the dinucleotide (n=16; AA, AT, CA, CG, etc.) frequencies at the start of each sequence (e.g. at *−*1500 from *µ*). These probabilities are used to initiate the sequence generator. The second distribution obtained from the sequence data is the per position conditional probabilities (e.g. *P* (*n_i_|n*_(_*i −* 1))) (see Supplemental Figure 4). The dinucleotide and conditional probabilities are described by equation 2. Position specific mononucleotide frequencies, which simply reflect the probabilities of a given nucleotide at a given position across the window, were calculated for Figure 1C. Position independent (also referred to as “flat”) probabilities (shown in Figure 1D, left) were generated by taking the mononucleotide probabilities and averaging them across the window (*H*=1500*∗*2).

Note that base composition plots (Supplemental Figure 4A and Figure 1C) are smoothed for clear visualization (scipy savgol filter version 1.5.4). Code used to calculate the position specific probabilities can be found within the sequence generator module of TF profiler.

### Generating simulated sequences around RNAPII initiation

Using the dinucleotide training data described in Section Calculating positional probabilities we employ a Markov chain to generate 10^6^ sequences each from the promoter and enhancer probability sets. This was achieved by using numpy (version 1.19.5) random number generator based on 1) the initial dinucleotide probability and 2) the subsequent conditional probabilities that account for position X-1 to select the nucleotide in position X. The sequences were checked to ensure there was no identical sequences within the 2*∗*10^6^ sequences generated. The validity of sequence generation was confirmed by ensuring that generated sequence recapitulate the probability distributions used in their generation (within *±* 0.0001). Sequences were generated in batches with distinct numpy seeds (seeds used: 38-50, 107-119, 275-287, 395-407, 462-474, 523-535, 687-699, 721-733, 831-843, 986-998) and the probabilities used are available on the additional data github page. The generation of mononucleotide and flat simulated sequences were generated in a similar manner (including the same seeds), only using the mononucleotide and position independent probabilities respectively. Code used to generate all sequences can be found within the sequence generator module of TF profiler.

### Counting over genes and bidirectionals

RefSeq gene counts for human sample within DBNascent were counted over hg38 RefSeq genes (hg38 release GCF 000001405.40-RS 2023 03) using Rsubread, featurecounts (version 2.12.3)[60, 76]. For all samples within a biological replicate for a given data set (both control, n=299 and perturbation, n=411 biological replicates), the mean RPKM was calculated for every gene isoform. Only the highest mean RPKM isoform for every gene was retained. Gene counts were used for additional analyses, including the transcription level of the gene encoding the TFs across tissue types, the transcription level of TF genes vs TF activity and DESeq2 analyses between control and perturbation conditions.

Bidirectional counts were also measured to assess 1) whether GC content of bidirectionals relates to the transcription level and 2) how this relates to enhancer and promoter content. This data is shown in Supplemental Figure 3. To count over all bidirectionals, the master bidirectional file was utilized (generation of this file described in Section Calculating positional probabilities). This file contains all bidirectionals called within the 126 control data sets. To ensure that the regions were wide enough, the regions within the master bed file were windowed +/-150bp surrounding *µ*. This could cause some regions to overlap, therefore the bedfile was sorted (sort -k1,1 -k2,2n) and merged (bedtools merge, version 2.28.0). Feature counts was used to count over the windowed master file using Rsubread, featurecounts (version 2.0.1). Like with genes, all individual control biological replicates (n=299 independent samples from n=126 control data sets) were used to count over the windowed-master bed file.

### Motif scanning

Motif scanning was performed using the MEME suite (version 5.0.3) function FIMO scan[77]. This scan was performed using a flat background model (equal distribution assumed of the four canonical nucleotides). The threshold was set to 1e-5. The motif files used were from HOCOMOCO version 11[39]. The scan was performed across the human genome (hg38) and these motif hits were used for subsequent analysis. Internal to the TF profiler program, the motif scan can also be performed de novo across only the bidirectional regions provided, or take in pre-scanned regions genome wide. Motif scanning was performed on simulated sequences using the same parameters. Code used to perform motif scanning can be found within the fimo scanner module of TF profiler.

### Calculating distances between RNAPII initiation and motif hits

To measure TF co-localization, the relative distance between a motif hit and the center of the bidirectional transcript must be assessed. The distance for all motif hits within the large window (*H*=1500) of a given region was calculated, using the center of the motif and the center of the bidirectional. For motifs and regions of odd length the center is rounded to the nearest even integer per the native python rounding function. Each motif hit is associated with the distance to the center of the bidirectional as well as two ranking metrics. The two ranking metrics are a distance rank metric (e.g. which motif is closest to the center of the bidirectional, where 1 is closest) and a quality rank (e.g. defined by FIMO score where 1 is the highest quality hit in the region). All motif hits within the large window are stored within the distance tables.

For this study only a single motif hit for a given PSSM per bidirectional was retained for further analysis. Hence, for each bidirectional region and *distinct* PSSM, only a single hit per *unique* PSSM is considered for further analysis. In the case of multiple motif hits for a single PSSM within one bidirectional, the motif hit used for further analysis was the highest quality motif hit (ie quality rank=1). If there were two (or more) motif instances with equivalent high quality rankings, the high quality motif hit closest to the center of the bidirectional was used (ie distance rank=1). Code used to generate these distance tables can be found within the distance module of TF profiler.

### PSSM GC content analyses

To calculate the GC content of the PSSMs we extracted all probabilities for both G and C across the length of the PSSM and summed them together. This was then divided this by the length of the PSSM to give the overall probability of a GC within the PSSM itself. This was done for all HOCOMOCO core TFs. To validate that the GC percentage is associated with a given TF rather than genomic context (nucleosome arrangement, for example), we looked at both SELEX and protein binding microarray data (CIS-BP version 2.00). The GC content was calculated for PBM and SELEX in the same manner as HOCOMOCO PSSMs (Supplemental Figure 8A-B).

### Calculating MD-scores

The calculation of MD-scores was originally defined in Azofeifa et. al. [21] and is described mathematically by equation 1. Briefly, the MD-score quantifies colocalization of motif instances (hits) near sites of RNAPII initiation (*h* = 150bps) relative to a larger local window (*H* = 1500bp) genome wide. The MD-score was calculated for all motifs within HOCOMOCO core version 11 (n=388 motifs)[39], in every data set in this study (n=287 data sets).

To calculate expected MD-score from simulated data, we leverage each data set’s distinct proportion of enhancer to promoter bidirectionals (Supplement Figure 5) – thus accounting for each data sets’ distinct composition profile. To this end, we calculate the proportion of promoter associated bidirectionals (see Section Curating a master bidirectional region list for labeling promoter bidirectionals). The proportion of promoter associated bidirectionals ranges from 0.14-0.49 across the 287 data sets. Based on this for every 0.02 step from 0.14-0.5 we calculated simulated MD-scores using 10^6^ simulated sequences (and associated motif hits) total. To do so we used numpy random number generator to randomly select a given proportion of promoters from the 10^6^ promoter sequences and the remainder from the 10^6^ enhancer sequences. For example, if a given data set had a proportion of 0.26 promoters, then 260,000 promoter sequences and 740,000 enhancer sequences were be selected from the dinucleotide simulated sequence data. From these 10^6^ sequences the expectation MD-scores were calculated and used for the basis of comparison for subsequent analyses. For each data set the MD-score proportion was rounded to the nearest 0.02 (a proportion of 0.255 rounds to 0.26; a proportion of 0.245 rounds to 0.24), and the expected MD-scores are selected from that set as the background model. Five seeds (96, 118, 559, 603, 961) were used for the numpy random number generator to subset the sequences. All resulted in similar expectation MD-scores for a given TF within a promoter proportion set. The seed used for selecting promoter and enhancer sequences for all subsequent analysis was 118.

### Statistically assessing TF profiles

We sought to statistically assess whether the MD-score for a given TF was higher (ON-UP) or lower (ON-DOWN) than expectation for each data set. For the meta-analysis, TFs across all data sets were combined for subsequent linear fits. Two separate fits were conducted, one on the control condition data Supplemental Figure 6A, n=48,888 points, 126 data sets with 388 TFs) and one on perturbation conditions (Supplemental Figure 11A, n=62,468 points, 161 data sets with 388 TFs).

In each case when all data is fit the slope is greater than 1, indicating higher activity in the experimental data than the expectation model. This is an expected result as some TFs should be ON and active in a given cellular context. Therefore we opted to use an inlier method, where we fit a set proportion of inlier TF MD-scores to a linear regression. The proportion of inliers was optimized for each set independently by testing every 5% inlier proportion from 5% (almost no TFs being fit) to 100% inliers (all of the data). The proportion closest to slope of 1.0 and intercept of 0.0 is assumed to identify the set of TFs unchanging within the set of data. The normal distribution of the residuals of the inliers was then used to attribute a p-value for each TF across all data sets (Supplemental Figure 6B, Supplemental Figure 11B). While the data was fit all together to get a better estimate the distribution of the residuals, the TF profiler program fits the residuals of the inliers for a single data set at a time by default for single case uses. Code used to generate these distance tables can be found within the statistics module of TF profiler.

### Clustering TF profiles

To generate the highest confidence TF profiles for a given TF we required a degree of replication across tissue types in control samples. For tissues with at least four data sets, a TF within the TF profile needed to be called as ON in at least 50% of the samples. For tissues with sparse data (less than four data sets) this replication was not required. There were a total of 26 tissue classifications for defining the high confidence TF profiles: blood (hematopoieticprogenitor), blood (K562), blood (lymphoid), blood (marrow), blood (myeloid), bone, brain, breast, embryo, eye, heart, intestine, kidney, liver, lung, lung (fetal), lung (muscle), muscle, ovary, prostate, skin, skin (foreskin), skin (lymph), stem cell, umbilical, uterus. Which data set belongs in which tissue set is defined in the sample metadata table found on in the additional data github. The tissues were defined in narrow categories as TFs vary between cell types as well as between tissues. The narrow classification permits for greater sensitivity when defining high confidence TF profiles.

Once tissue specific profiles were rigorously defined they were clustered using the R (version 3.6.0) package ComplexHeatmaps (version 2.2.0) which utilizes the native R function hclust. For clustering purposes the TF profiles were numerically represented as 1 (ON-UP), −1 (ON-DOWN) and 0 (OFF). The Euclidean distances were used followed by Wards method to cluster the profiles. A full cluster map was generated and shown in Supplemental Figure 7. For the main text figure the tissues were manually divided into three categories, tissue, organ and developmental. The tissues within those categories were clustered to assess which are most closely related. This ordering was used for Figure 2B.

### Classifying TFs

Here we define three categories of TFs: ubiquitous, tissue specific and stimulus responsive. All classifications were defined using the high confidence TF profiles. High confidence TF profile generation is described in section Clustering TF profiles.

To classify a TF as ubiquitous is was required to be ON-UP in at least 95% of the control data sets. To classify a TF as tissue specific, it needed to be uniquely ON in a given tissue. The exception for this is blood TFs (such as GATA and STAT TFs). These TFs had strong signatures in blood but also tended to be called ON in many organ samples. For this reason, blood specific TFs were excluded from being called ON in developmental or organ sets. Additionally, many organ TFs were shared due to similar function across tissues. If a TF in organ samples was only shared across two organs it was still defined as tissue specific. One category not discussed in depth is shared, but not ubiquitously shared TFs. This general group is classified as TFs that are on in more than two organs, more than one blood cell type or more than one developmental cell type. Finally, the stimulus responsive TFs defined by 1) TFs that were called ON in the perturbation sample but not called ON in the control sample and 2) not a ubiquitous or tissue specific TF within the tissue of the experiment tested. TF classifications are outlined in the TF classes table found on in the additional data github.

### Comparing bidirectional regions across tissue types

To compare region usage in control conditions, each tissue was assigned a consensus region set. To generate consensus regions across tissues the master bed file (described in section Curating a master bidirectional region list) was used. For each region within the master bed file, the data set that contributed to that region was noted. The tissues were broken into broad categories (n=15; brain, blood, muscle, fetal, liver, ovary, hematopoietic, breast, skin, kidney, eye, umbilical, bone, prostate, and uterus) for region selection to increase the total number of regions accounted for in subsequent comparisons. Which data set belongs in which broad tissue set is defined in the sample metadata table. For a region to be called within the consensus profile, the region needed to be attributed to at least 50% of the data sets within the broad tissue set. This ensures that the regions called are truly active bidirectional regions within the broad tissue category. Distances between the consensus regions and all HOCOMOCO motif hits were calculated using the distance calculations described in section Calculating distances between RNAPII initiation and motif hits. Motif hits within *±*150bp of initiation for a given bidirectional were considered a positive hit. Positive motif hits and consensus regions were used to systematically assess TF region selection across tissues, as shown in Figure 3C.

The perturbation condition TNF*α* is one of the most highly represented perturbations, with 6 data sets in 6 independent publications applied across a myriad of tissue types. To study region selection across tissue type in TNF*α* conditions we used *muMerge* across the 6 independent TNF*α* data sets to create a master TNF*α* bed file. Contributions to each region per data set were retained. These regions were used for distance calculations with all HOCOMOCO motif hits. As previously, motif hits within *±*150bp of initiation for a given bidirectional were considered a positive hit. This data was used to assess NF*κ*B subunit region selection as shown in Figure 5C and Supplemental Figure 14.

### Using ChIP-seq data from CistromeDB

Both TF and histone ChIP-seq region data was obtained from CistromeDB[61, 62]. Within this database there are six total quality parameters assessed for every ChIP-seq experiment. These can be broken into two main categories, mapping and peak quality. The mapping scores account for sequence quality, number of unique sequences and unique molecule representation after sub-sampling the data. The peak scores account for the number of peaks, the signal to noise ratio and the overlap of peaks with accessible regions. In order for the ChIP-seq sample to be used here, we required the sample to pass at least one parameter within both mapping and peak scores. CistromeDB contained TF ChIP-seq data for 316 unique TFs within HOCOMOCO v11 core set (n=388 total) that passed the defined QC standards.

Promoter associated ChIP-sites were defined as a ChIP site falling within 1000bp (300bp upstream, 700bp downstream) of RefSeq (hg38 release GCF (109.20190607 2019 06) annotated transcription start site (TSS), as with bidirectional calls. The percentage of promoter-associated regions was calculated by the total number of promoter associated ChIP-sites over the total number of ChIP-sites that fall within a bidirectional region from the master bed file. This reduces noise and regions where a TF is bound but not actively regulating. Thus the ChIP promoter percentage reflects the percentage of functional TF binding events that occur within promoter regions versus all functional binding events.

In many cases, there were multiple ChIP-seq samples for a single TF. In this case, the median calculated promoter percentage was used. Heat maps in Figure 1 use cistromeDB regions from five independent samples per condition detailed in the ChIP metadata table (found on the additional data github page). Distance tables were generated using the TF profiler program as previously described in Section Calculating distances between RNAPII initiation and motif hits. The R (3.6.0) package ComplexHeatmaps (version 2.2.0) was used to plot the motif localization using the generated distance tables.

### Fit classification for transcription level of TFs

We construct a simple classifier to assess the distribution of the transcription level for all genes encoding TFs across all data sets. To do this we used Fitter (version 1.5.2) built on scipy (version 1.5.4). This program takes an array RPKM normalized counts across control data sets (as described in section Counting over genes and bidirectionals) and assesses how well that array fits a given data distribution. We used Fitter to classify the transcription level distributions as either exponential or normal, with the best fit defined as the minimum sum of the error metric squared (parameter fitterf.get best(method = ‘sumsquare error’)). If the KS p-value was greater than 0.1 this indicated a poor fit for both exponential and normal, thus the TF was classified as “other.” (Supplemental Figure 9).

We generated the cumulative frequency plots for the TF gene transcription across control samples using a bin size of 0.001 RPKM. The mean cumulative frequency across high confidence TFs was plotted as a black line and the standard error is shown as a colored region across the high confidence TFs (Figure 4B). The high confidence TFs were selected by significance values (p-value *≤* 0.001). These TFs were parsed down to a group of TFs with a similar range of transcription level such that they could be plotted on the same x-axis for the cumulative frequency plots. The most confident tissue specific factors within their respective tissue types were identified as MyoD, GRHL2, TEAD4, p63, GATA1 and Oct4. The most confident ubiquitous factors across all tissue types were identified as NFYB, ELK1, CREB1, SP1, ATF1 and SP3.

### Single cell RNA-seq data

For single cell RNA-seq data (scRNA-seq) we used data published from the “human cell atlas of fetal gene expression”[63]. This data was accessed from NCBI GEO accession number GSE156793. We used the publicly available file titled: “GSE156793 S6 gene expression celltype.” To be considered “expressed” in a given tissue we used the expression cutoff of 0.01. scRNA-seq expression values were fit to either an exponential or normal distribution by Fitter as described in Fit classification for transcription level of TFs.

### Differential expression

Gene counts previously quantified from DBNascent were used for differential expression analysis (see section Counting over genes and bidirectionals). We focused on 6 data sets in which there was TNF*α* treatment and their corresponding controls. Data sets from the same tissue type were grouped within a single DESeq2 (version 1.26.0) object. Differential gene expression was assessed between TNF*α* vs control separately for the 4 tissues represented (heart[26, 66] n=8 control samples, n=7 treatment samples; lung[68, 69] n=5 control samples, n=5 treatment samples; intestine[67], n=2 control samples, n=2 treatment samples; prostate[70], n=2 control samples, n=2 treatment samples). Additional sample information is defined within sample metadata table. Both the sample metadata table and differential expression results can found on the additional data github page. NF*κ*B targets were defined from the GSEA Hallmarks (version 5.0) pathway: TNF*α* signalling via NF*κ*B (n=200 genes).

## Funding

This work was funded by the National Science Foundation under grants ABI1759949 and the National Institutes of Health grant GM125871 and HL156475.

## Competing interests

Drs. Dowell and Allen are on a patent that uses enhancer RNAs to infer transcription factor activity. The other authors declare that they have no competing interests.

## Author’s contributions

TJ, MAA and RDD conceptualized the body of work and interpreted results. MAA and RDD supervised the study. DJT aided with conceptualization and interpretation. RFS and LS collected and derived quality control metrics for the nascent sequencing data within the DBNascent construction. TJ conducted all data analysis. MAA aided in code development. TJ and RDD wrote the paper. All authors revised the final manuscript.

## Acknowledgements

We are grateful to the BioFrontiers IT department for their support in building the database.

---

**Supplementary Figure 1.**
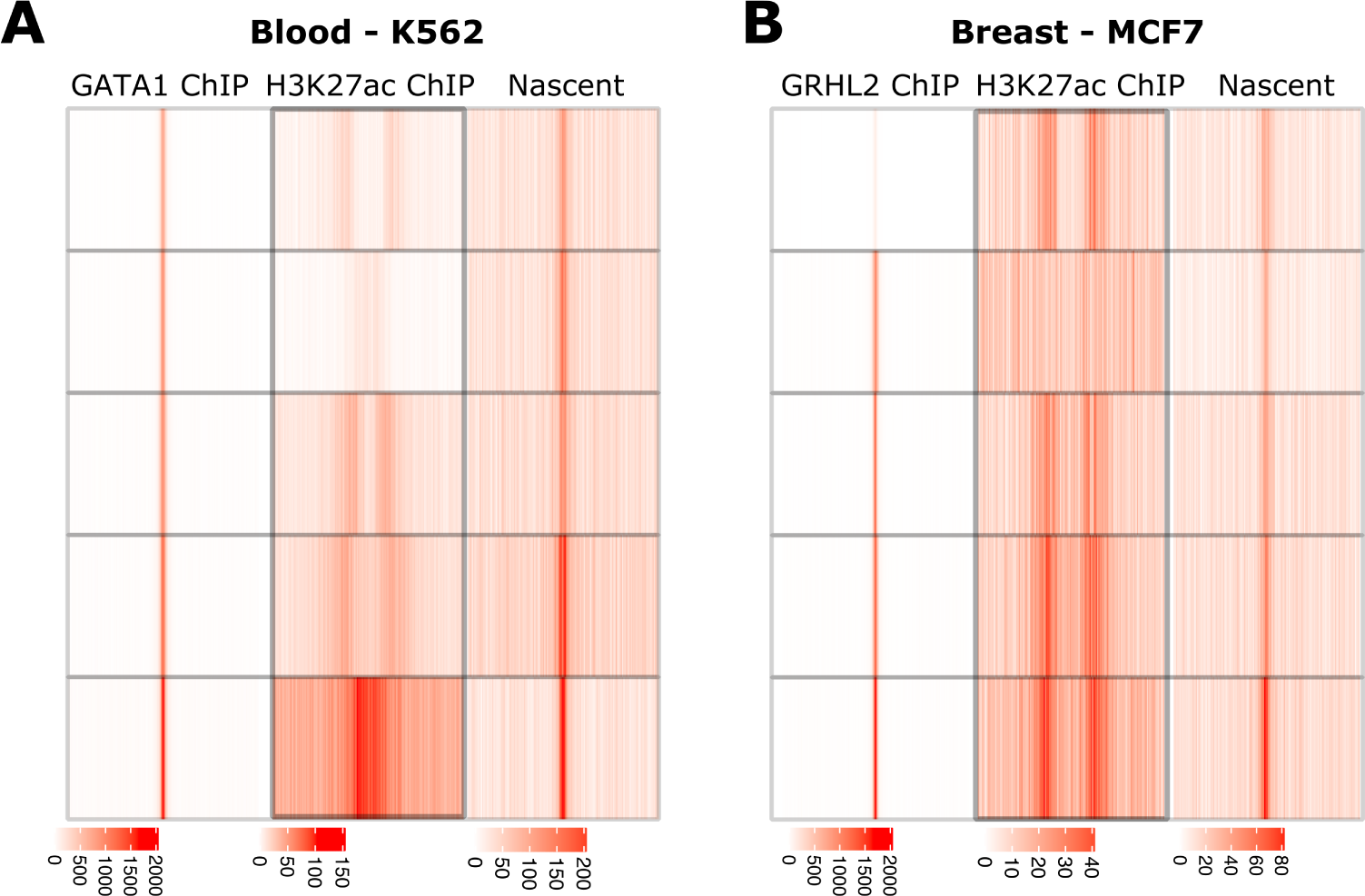
MD-scores calculated from ChIP-seq data compared to nascent PolII initiation data. Heat maps showing the co-localization of motif instances (hits to the PSSM) across three data types (TF ChIP left, H3K27ac middle, and nascent data right) for A: GATA1 in blood (K562) and B: GRHL2 in breast (MCF7) samples. In each case, five independent experiments were chosen for each example (top to bottom). ChIP-seq data was pulled from cistromeBD[61, 62]. Using panel A as an example, GATA1 ChIP-seq (left) demonstrates high co-localization of GATA1 binding with the GATA1 motif, as expected since ChIP-seq is how the GATA1 motif was defined. This is a low-throughput assay for annotating enhancers, as only one epitope can be pulled down at a time. H3K27ac ChIP-seq (middle) is a high-throughput assay to annotate enhancers. However, the broad histone peaks lose precision on the GATA1 motif:enhancer co-localization. Precision is required for robust MD-Score calculation[27, 78]. Finally, PolII initiation sites annotated from nascent sequencing data (right) demonstrate how a high-throughput assay can be used to precisely capture GATA1 motif:PolII initiation co-localization for robust MD-Score calculation. Importantly, the nascent transcription can be used to co-localize any TFs for which a PSSM is available. Similar results are seen with other TFs in other tissues (panel B: GRHL2 in breast tissue).

**Supplementary Figure 2.**
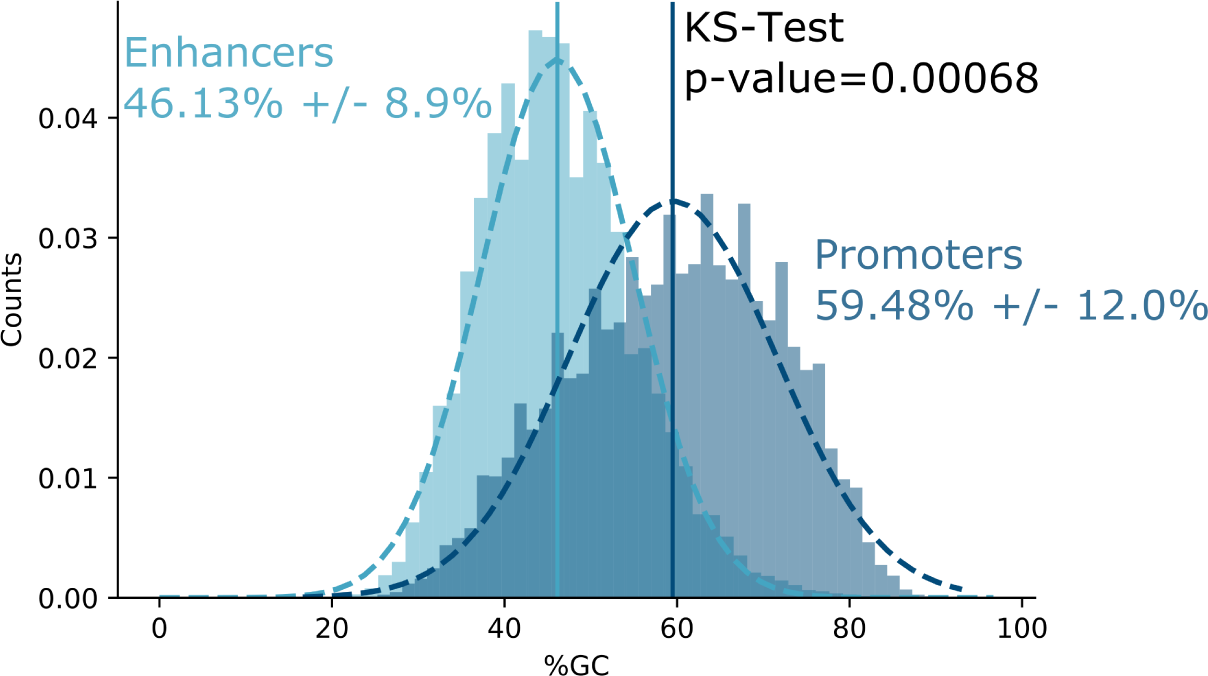
Conditional probabilities obtained from data. Distribution of PolII initiation region’s GC content follows a Normal distribution. Promoters are shown in dark blue with a mean GC content of 59.48%. Enhancers (defined as all bidirectionals that are not at annotated promoters) are shown in light blue with a mean GC content of 48.13%. The normal distributions between enhancers and promoters are statistically different (KS-Test, p-value=0.00068).

**Supplementary Figure 3.**
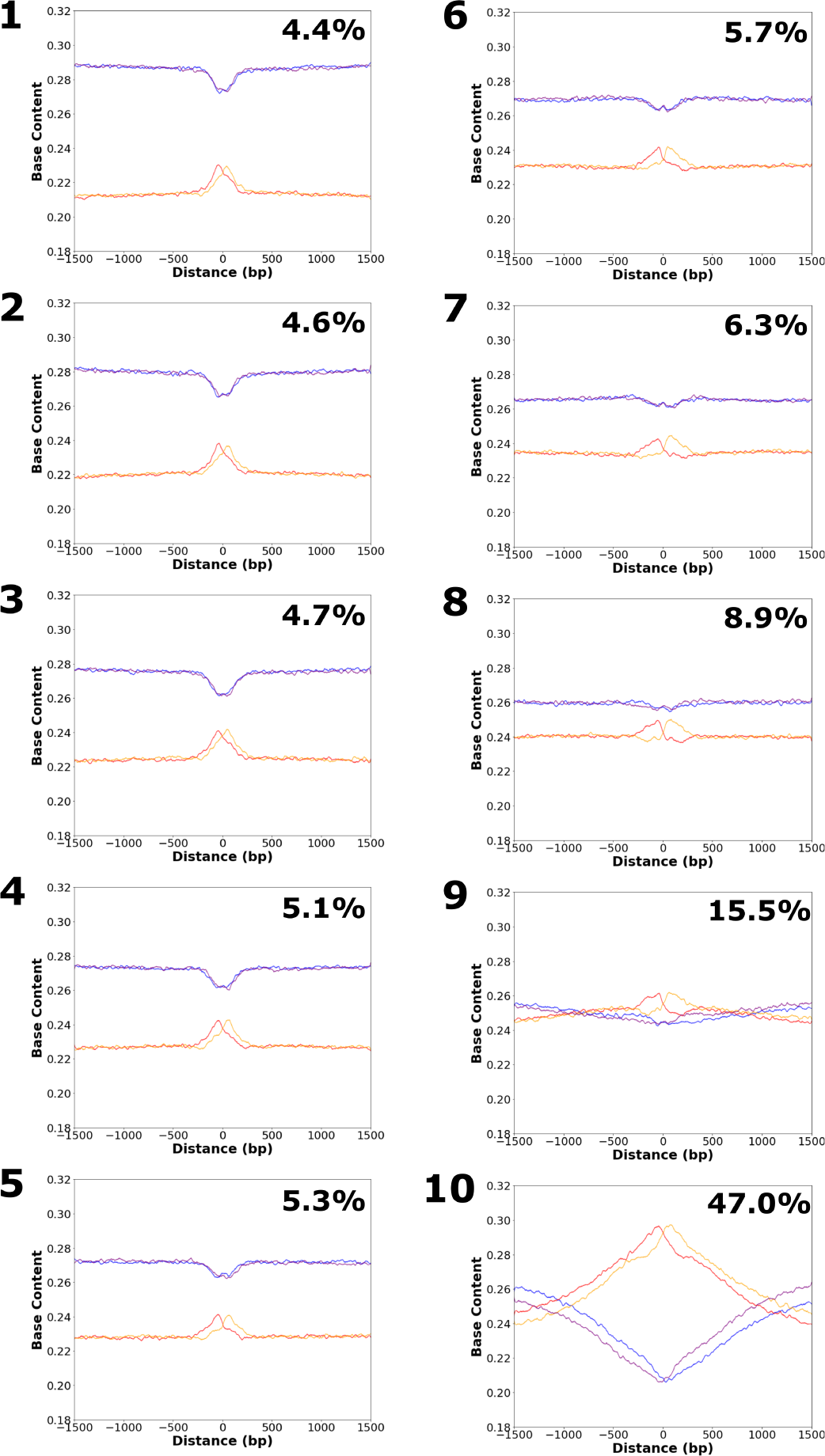
The highest transcription levels have the highest GC content. The 353,370 bidirectional regions identified in human breast samples were split into 10 bins ranked from the lowest transcriptional level (1) to highest transcription level (10). The base composition for each bin is displayed, showing increasing GC content with higher transcription level. For each bin the percentage of promoters represented within the bin is displayed in the top right corner of each plot. The total representation of promoters (by our definition of a bidirectional falling within −300bp/+700bp of an annotated TSS in Refseq) within these 353,370 regions is 10.8%. For low transcription bins (bins 1-8), there are few promoters and the AT content is higher than the GC content across the +/-1500bp window. The high transcription bins (9 and 10) have a higher promoter content and the GC content is greater than the AT content near the initiation region (*µ*).

**Supplementary Figure 4.**
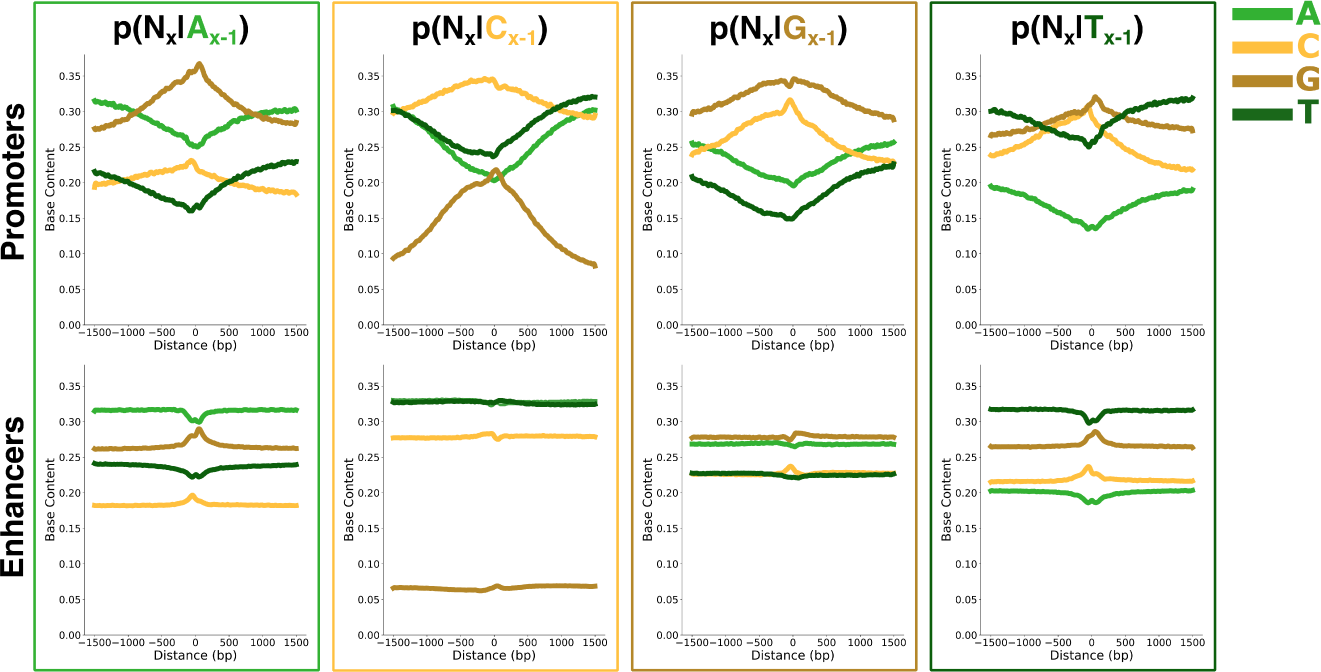
Conditional probabilities obtained from data. The conditional probabilities obtained for the background model. The probabilities derived from promoters (top) show stronger biases in conditional probabilities than those derived from enhancers (bottom). Each panel (from left to right A, C, G then T) shows the probability of the following base being a A (light green), C (light gold), G (dark gold) or T (dark green) given the base indicated at the top. Lines were smoothed for clear visualization (scipy savgol filter version 1.5.4).

**Supplementary Figure 5.**
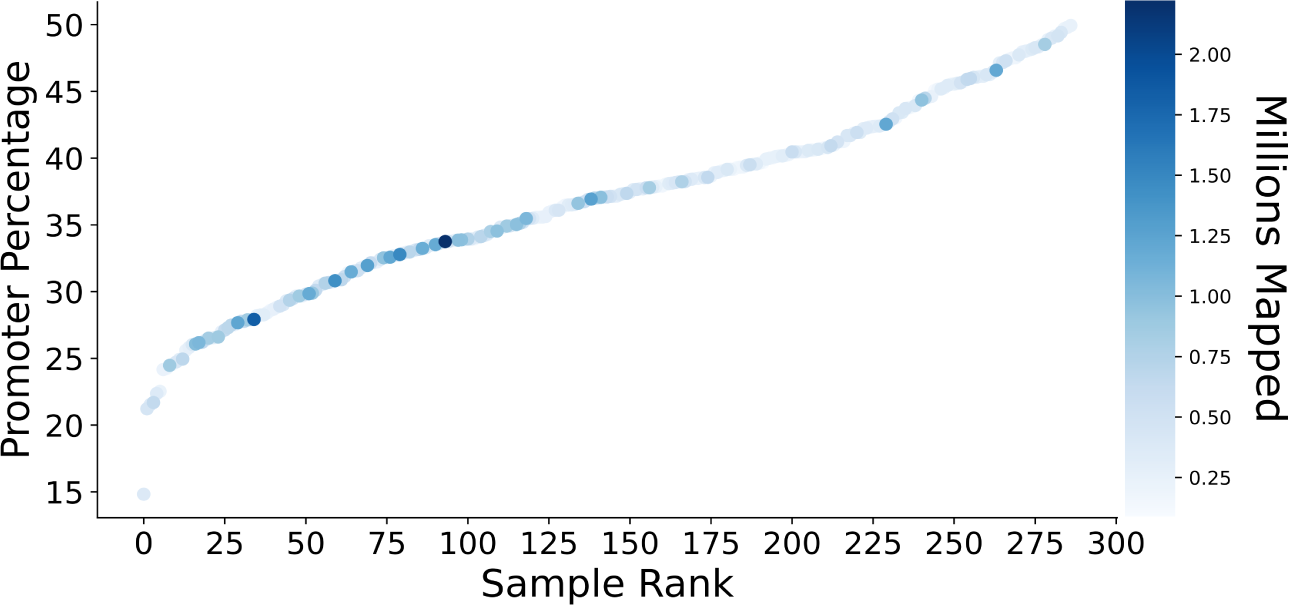
Representation of promoter percentage across 287 high quality samples. In this study 287 samples (representing unique cell types and/or treatment conditions across 89 publications) are assessed for the percentage of bidirectional peaks that fall within annotated promoters, colored by depth of data (millions mapped). As most enhancer bidirectionals are lowly transcribed, there is a bias for more deeply sequenced samples to have a higher percentage of enhancers and lower percentage of promoters relative to less well sequenced samples. The background for each individual sample is calculated using a promoter percentage equivalent to the observed value plotted here.

**Supplementary Figure 6.**
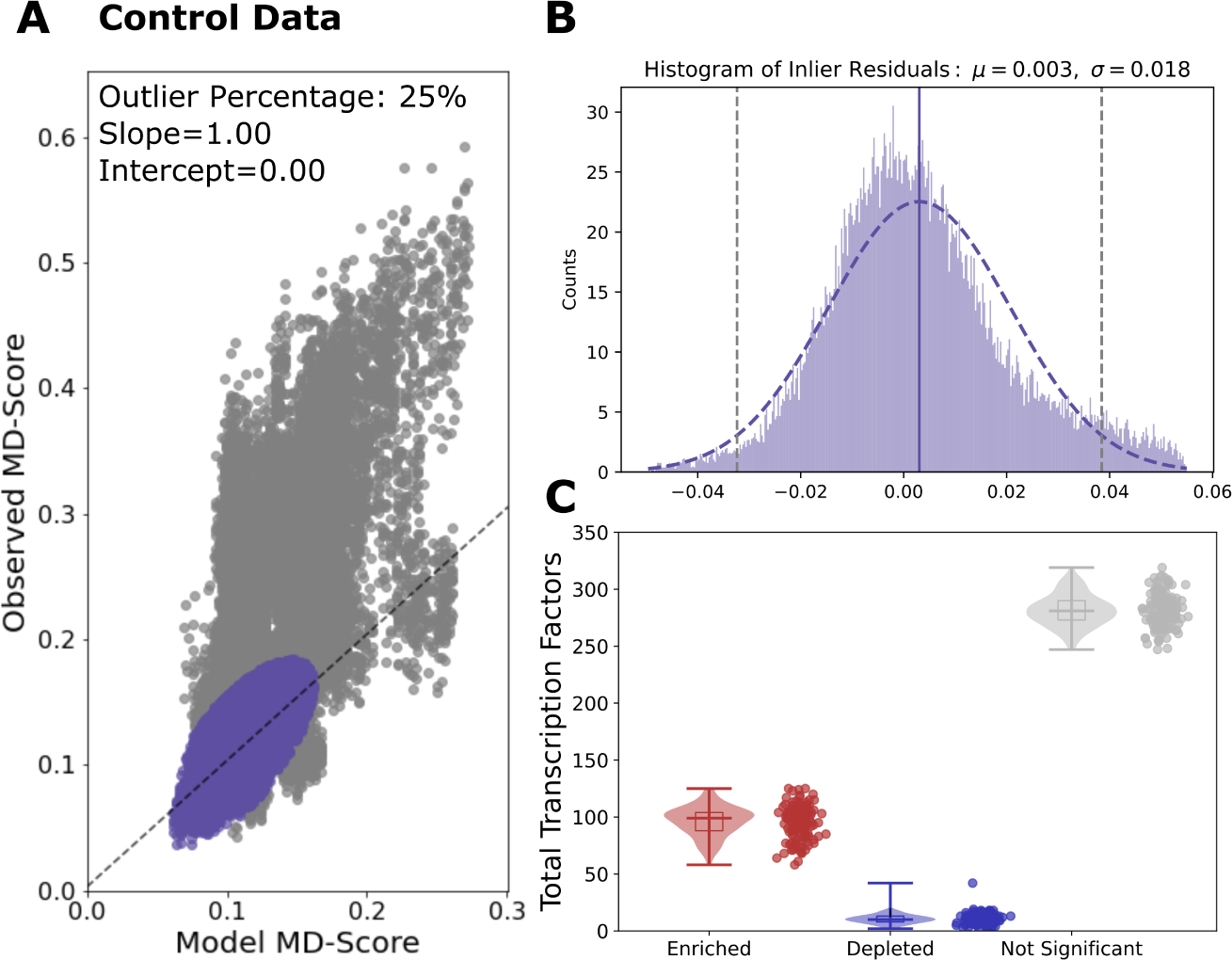
Statistical test of ratio of observed to expected MD-score in control conditions. A) The expected MD-score (x-axis) from the dinucleotide model and the experimentally observed MD-score (y-axis) were calculated for all HOCOMOCO TFs (n=388) from each control data set (n=126). For the set of all TF-data set pairs (n=48,888), the 75% inliers (purple) were fit to a linear equation resulting in a slope of 1.00 and an intercept of 0.00. The 25% outliers (grey) were excluded from the linear fit. B) The residuals of the inliers were fit to a normal distribution (purple) with *µ*=0.003 (dashed purple line) and *σ*=0.018 (2*σ* at dashed grey lines). These values were used to calculate significance on every TF within the control data sets (n=48,888). C) Violin plots demonstrating the number of TFs that are significantly enriched (ON-UP; red), depleted (ON-DOWN; blue) or OFF (grey) per data set within control conditions.

**Supplementary Figure 7.**
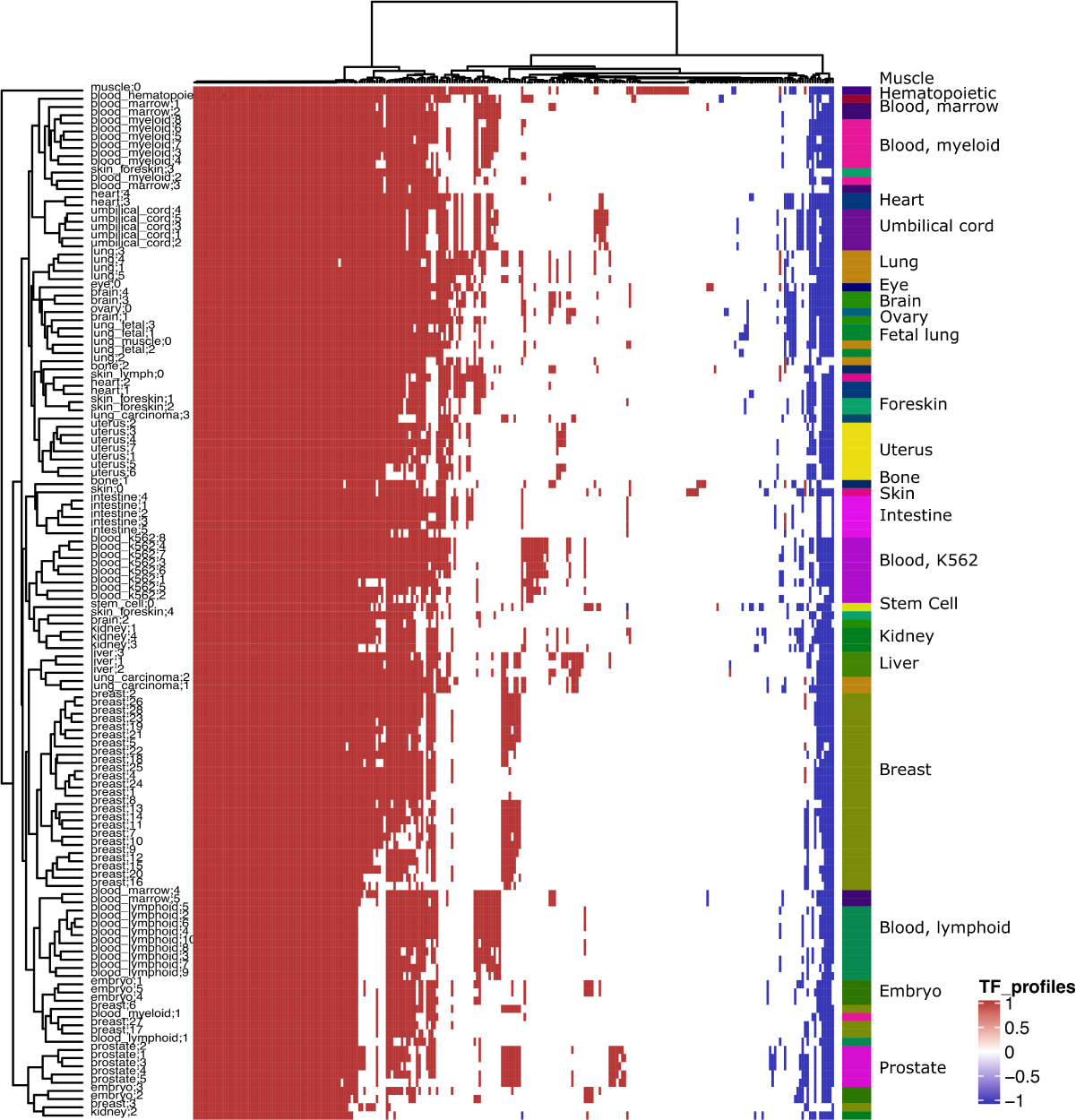
Control two dimensional cluster map. Related to Figure 2B, this is the cluster map for 126 TF profiles within control samples. This map is clustered in both dimensions using Ward clustering on euclidean distances. The primary driver of clustering is cell lineage.

**Supplementary Figure 8.**
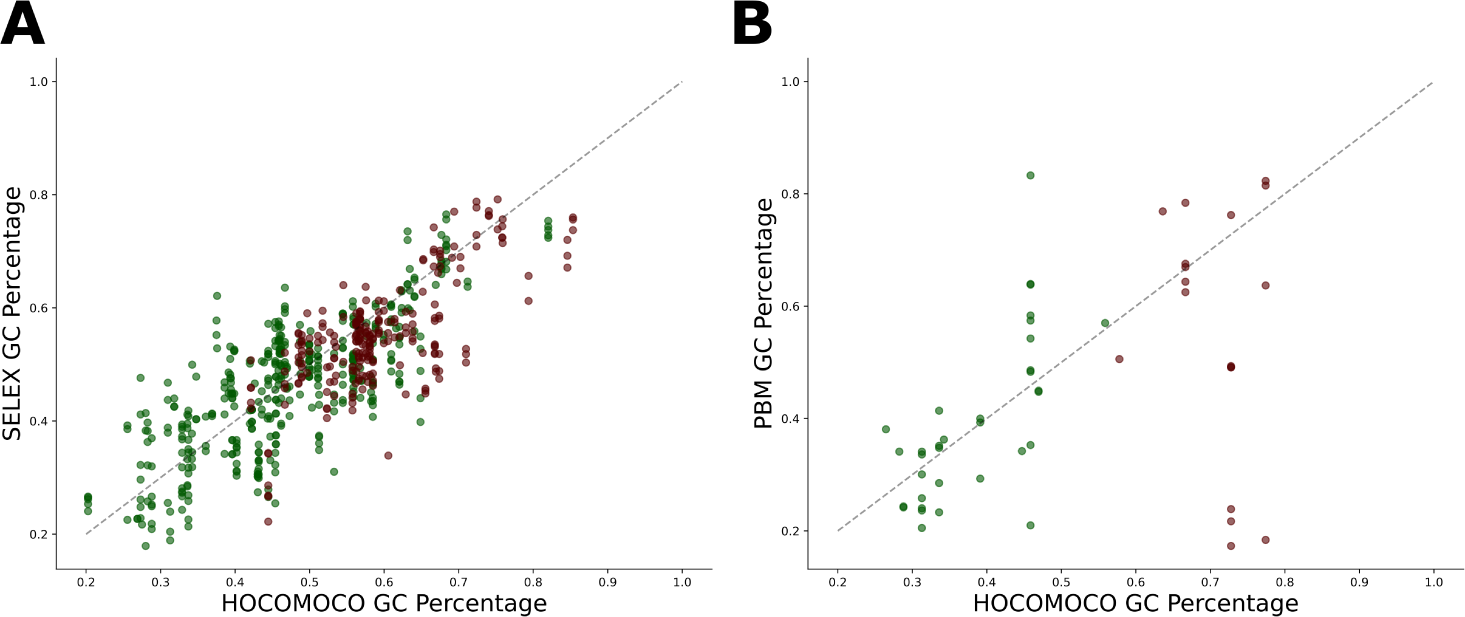
Ubiquitous TFs prefer binding more GC rich sequences compared to tissue specific TFs independent of genomic context. Scatter plot showing the calculated GC content from the HOCOMOCO PSSM (x-axis). As most PSSMs are derived from ChIP-seq data, we examined A) SELEX defined PSSMs (200 TFs, 915 SELEX defined PSSMs, y-axis) or B) Protein binding microarray defined PSSMs (20 TFs, 61 PBM defined PSSMs, y-axis). Tissue specific TFs are colored green, centered on the bottom left of the plot. Ubiquitous TFs are colored red, centered on the top right of the plot. The dashed line on each plot is the 1:1 line, which the points follow closely. There is a positive correlation between the HOCOMOCO PSSM GC percentage and the context independent derived PSSM GC percentage.

**Supplementary Figure 9.**
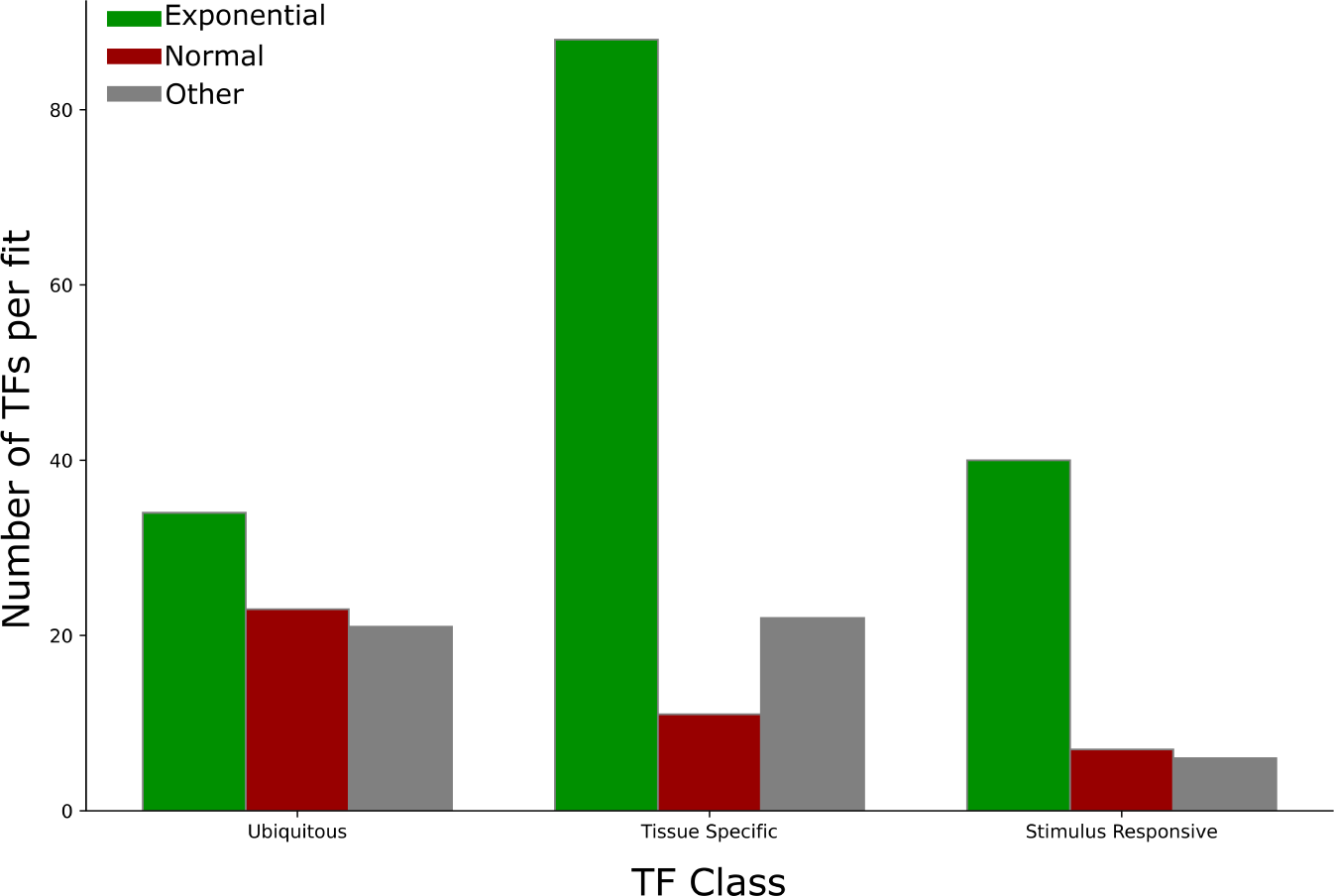
Transcription of the gene encoding the TF. Classification of the shape of the distribution of transcription for the gene encoding a TF. Bar plot showing the number of TFs per class that fit an exponential (green), normal (red) or other (grey) distribution. Other is defined by poor fit to both normal and exponential distributions (p-value $¿ 0.1). Proportionally, ubiquitous TFs fit a normal distribution more often than the other classes, indicating they are transcribed in all samples at a variety of levels.

**Supplementary Figure 10.**
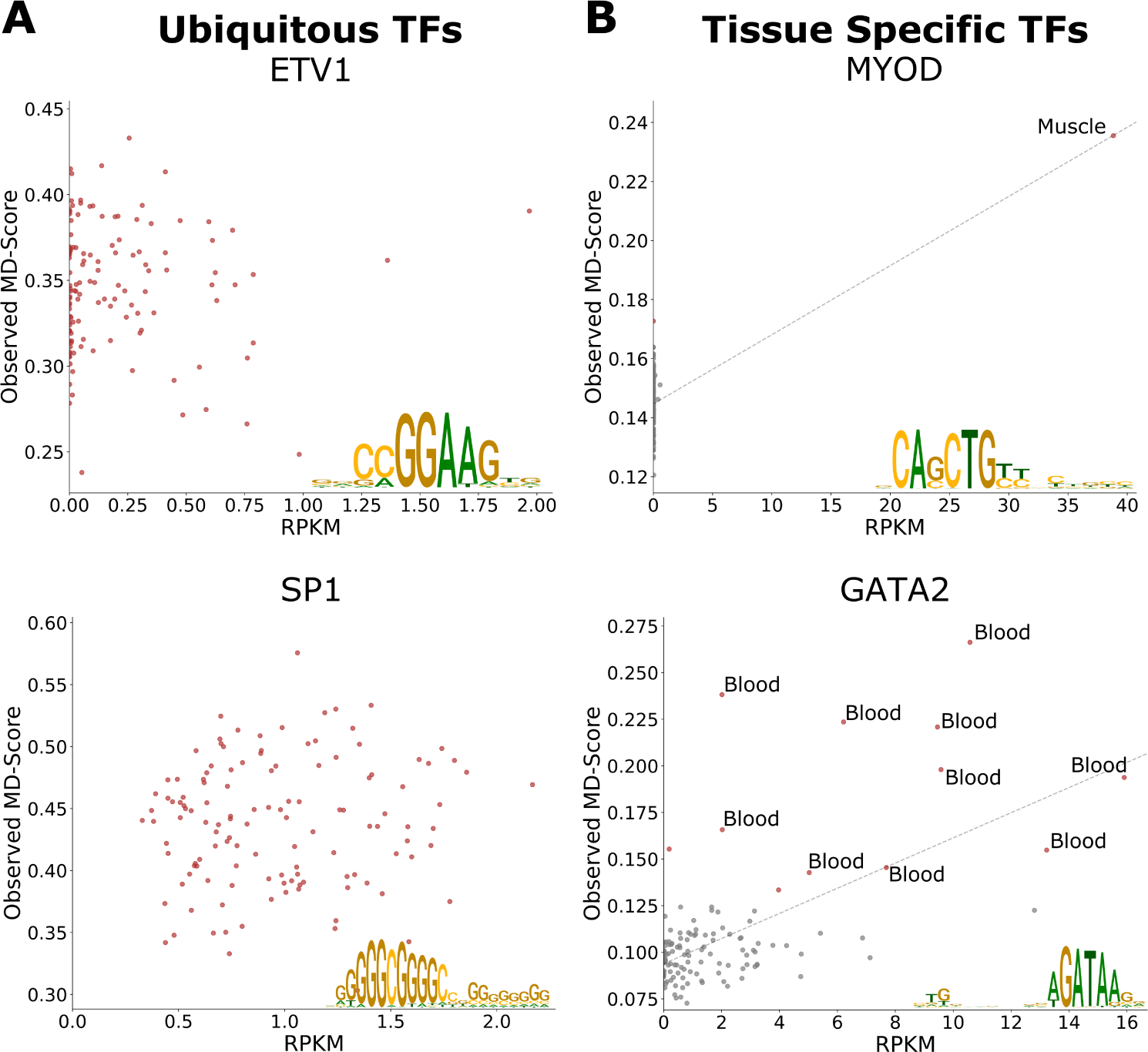
Additional scatter plots comparing transcription of the gene encoding the TF to the MD-score for the TF. Plots similar to Figure 4B-C. A) Two ubiquitous TFs: ETV1 (top) and SP1 (bottom). Inset are their respective motifs. Both show no correlation between transcription of the TF (x-axis) and its activity (y-axis). B) Two tissue specific TFs: MyoD (top; muscle specific) and GATA2 (bottom; blood specific). Inset are their respective motifs. Both show positive correlation between transcription of the TF (x-axis) and TF activity (y-axis).

**Supplementary Figure 11.**
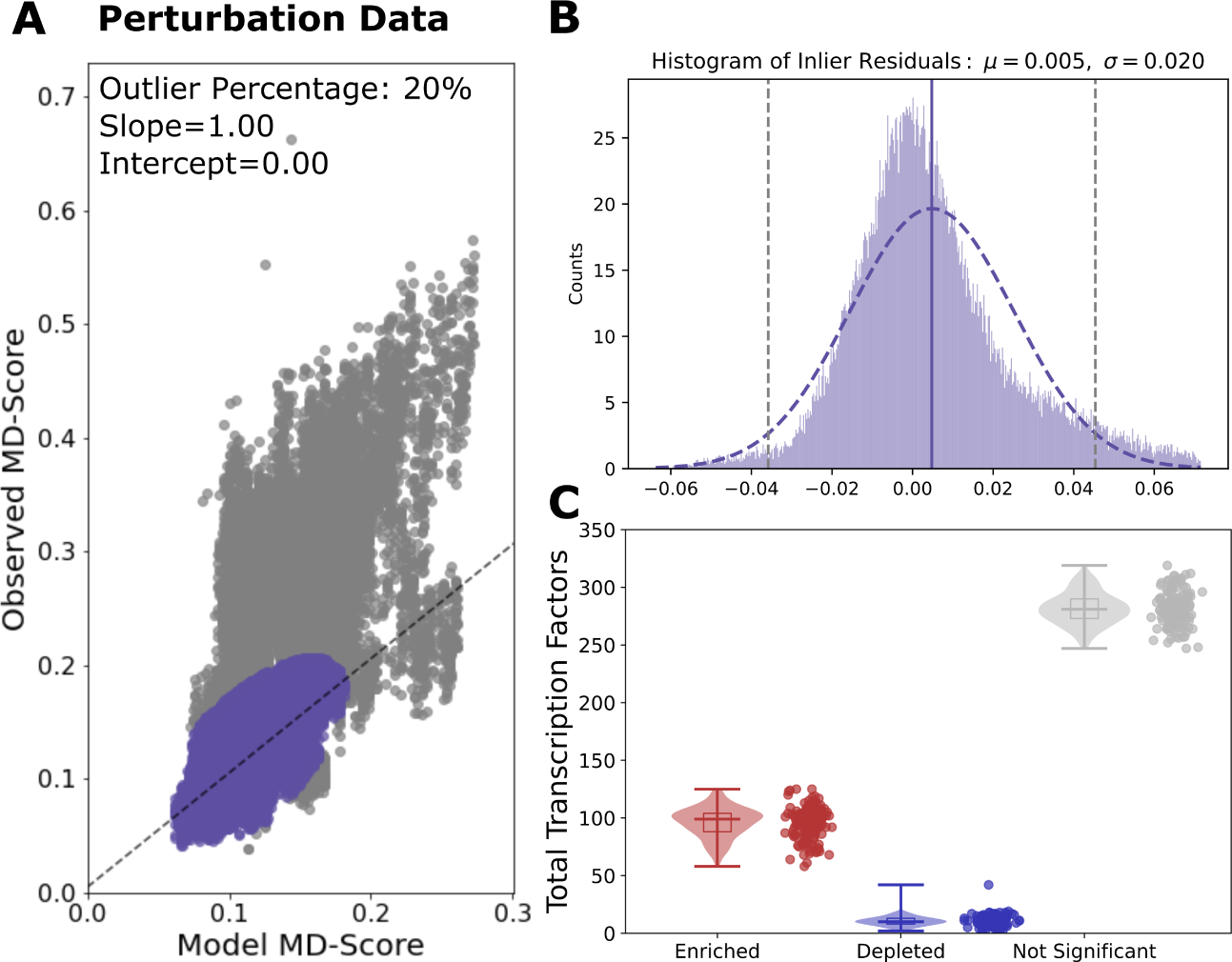
Statistical test of ratio of observed to expected MD-score in perturbation conditions. Figure similar to Supplemental Figure 6. A) The expected MD-score (x-axis) from the dinucleotide model and the experimentally observed MD-score (y-axis) were calculated for all HOCOMOCO TFs (n=388) from each perturbation or genetically modified data set (n=161). For the set of all TF-data set pairs (n=62,468), the 80% inliers (purple) were fit to a linear equation resulting in a slope of 1.00 and an intercept of 0.00. The 20% outliers (grey) were excluded from the linear fit. B) The residuals of the inliers were fit to a normal distribution (purple) with *µ*=0.005 (dashed purple line) and *σ*=0.020 (2*σ* at dashed grey lines). These values were used to calculate significance on every TF within the control data sets (n=62,468). C) Violin plots demonstrating the number of TFs that are significantly enriched (ON-UP; red), depleted (ON-DOWN; blue) or OFF (grey) per data set within perturbation conditions.

**Supplementary Figure 12.**
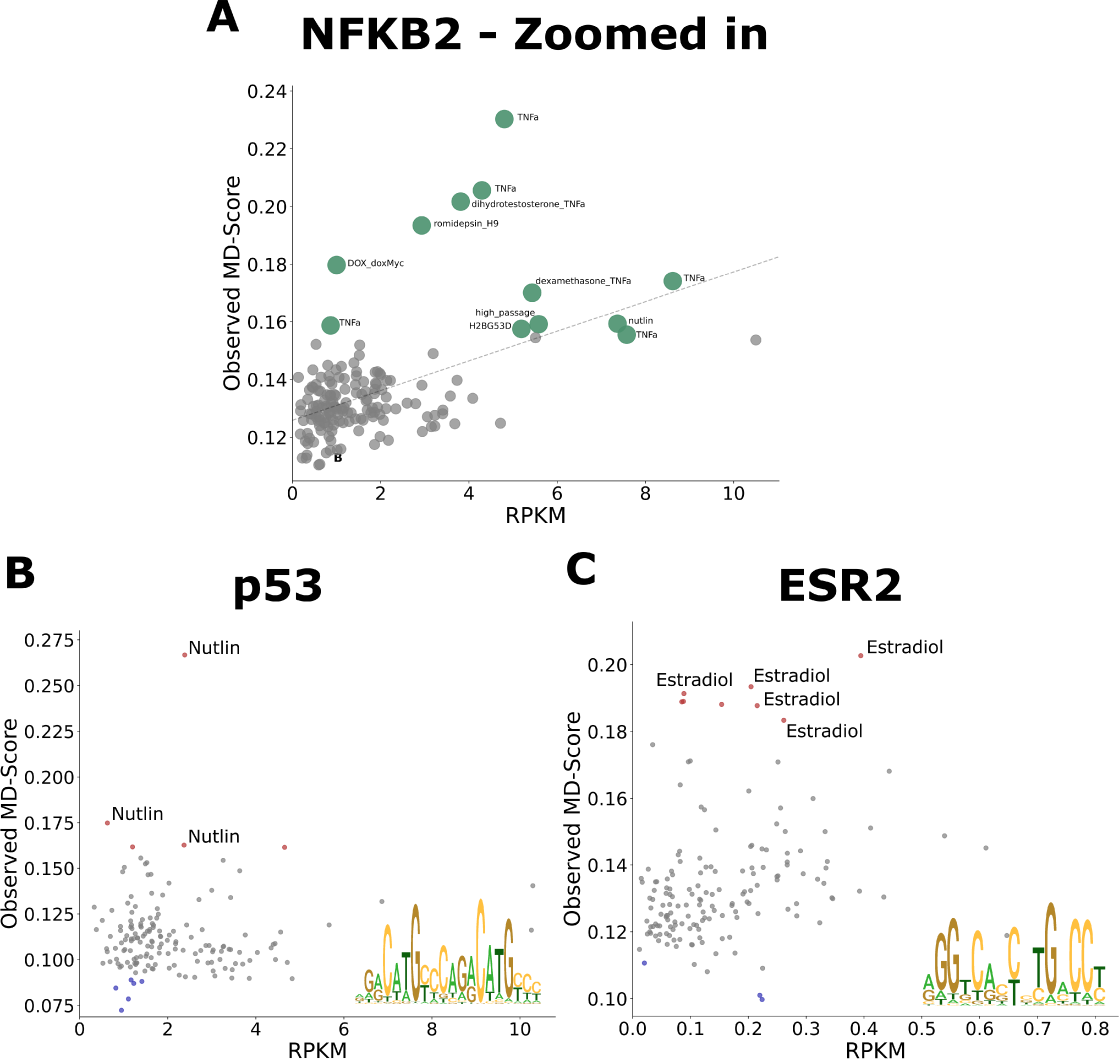
Additional scatter plots assessing stimulus responsive TFs. Scatter plots demonstrating the relationship between the transcription level of the gene encoding the TF (RPKM, x-axis) and MD-score (y-axis) for stimulus responsive TFs. A) A zoom in on the lower left of Figure 5B. Two examples: B) p53 and C) estrogen receptor. Graphs are similar to Figure 4B-C and Supplemental Figure 10. Inset are their respective motifs.

**Supplementary Figure 13.**
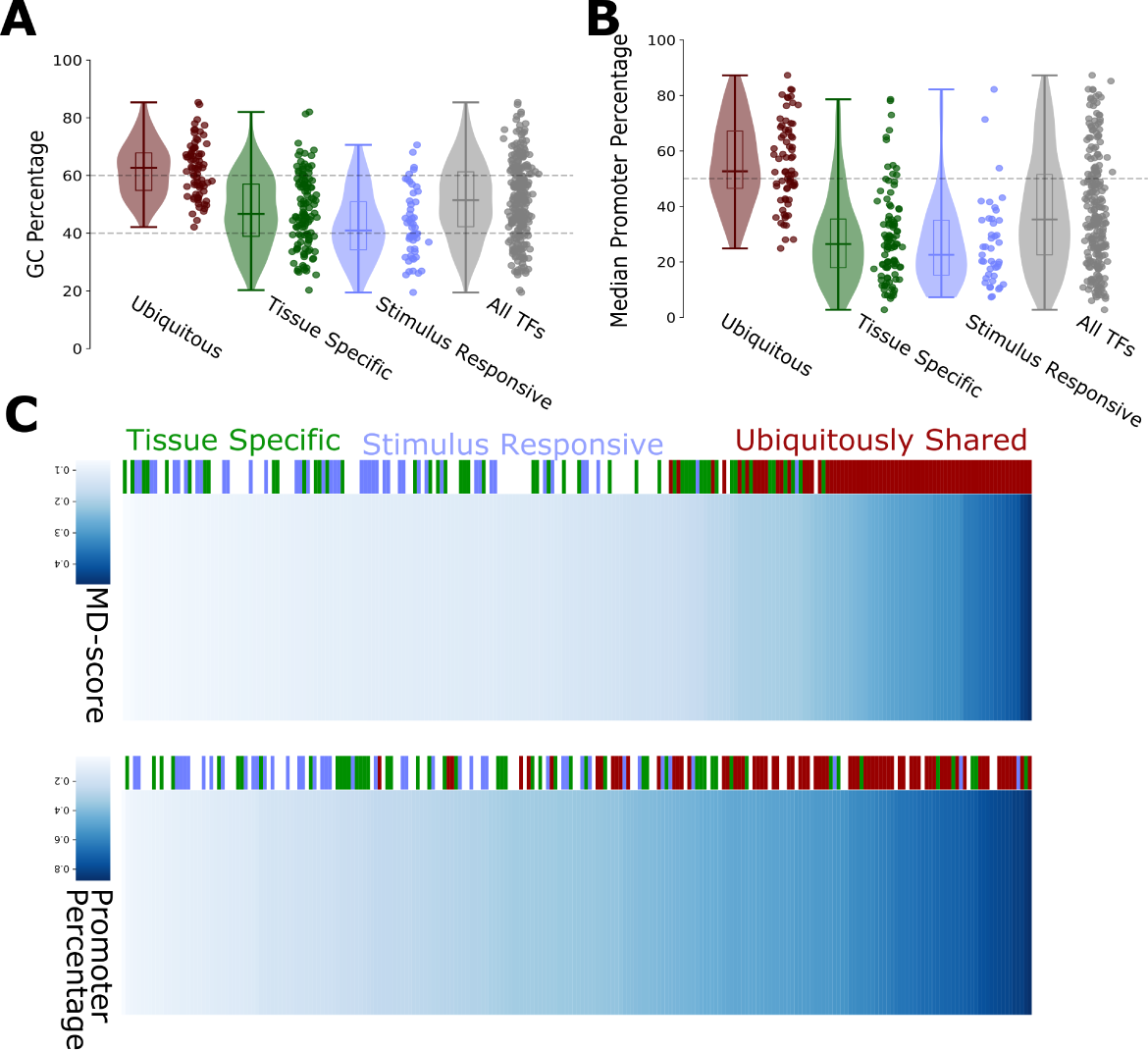
Stimulus responsive TFs display similar binding preferences to tissue specific TFs. A) Violin plots showing the GC content of the TF motif, similar to Figure 3A. We find that ubiquitous factors have motif biases close to that of promoters, whereas tissue specific and stimulus responsive factors have preferences closer to genomic background. B) Violin plot showing the median percentage of TF ChIP-seq peaks that fall within a promoter over peaks that fall within any bidirectional. We find that for ubiquitous factors that the TF binds within promoters more than half of the time (dashed horizontal line), whereas tissue specific and stimulus responsive bind at promoters less than 35% of the time (the median promoter binding percentage for all TFs). Ubiquitous (red), tissue specific (green), stimulus responsive (blue), and all TFs (grey). C) Similar to Figure 3D-E. In terms of both MD-score (top) an promoter percentage (bottom) stimulus specific TFs show a similar trends to tissue specific TFs and differ from ubiquitously shared TFs. In summary, stimulus specific TFs tend to have lower MD-scores on average and bind primarily at enhancers.

**Supplementary Figure 14.**
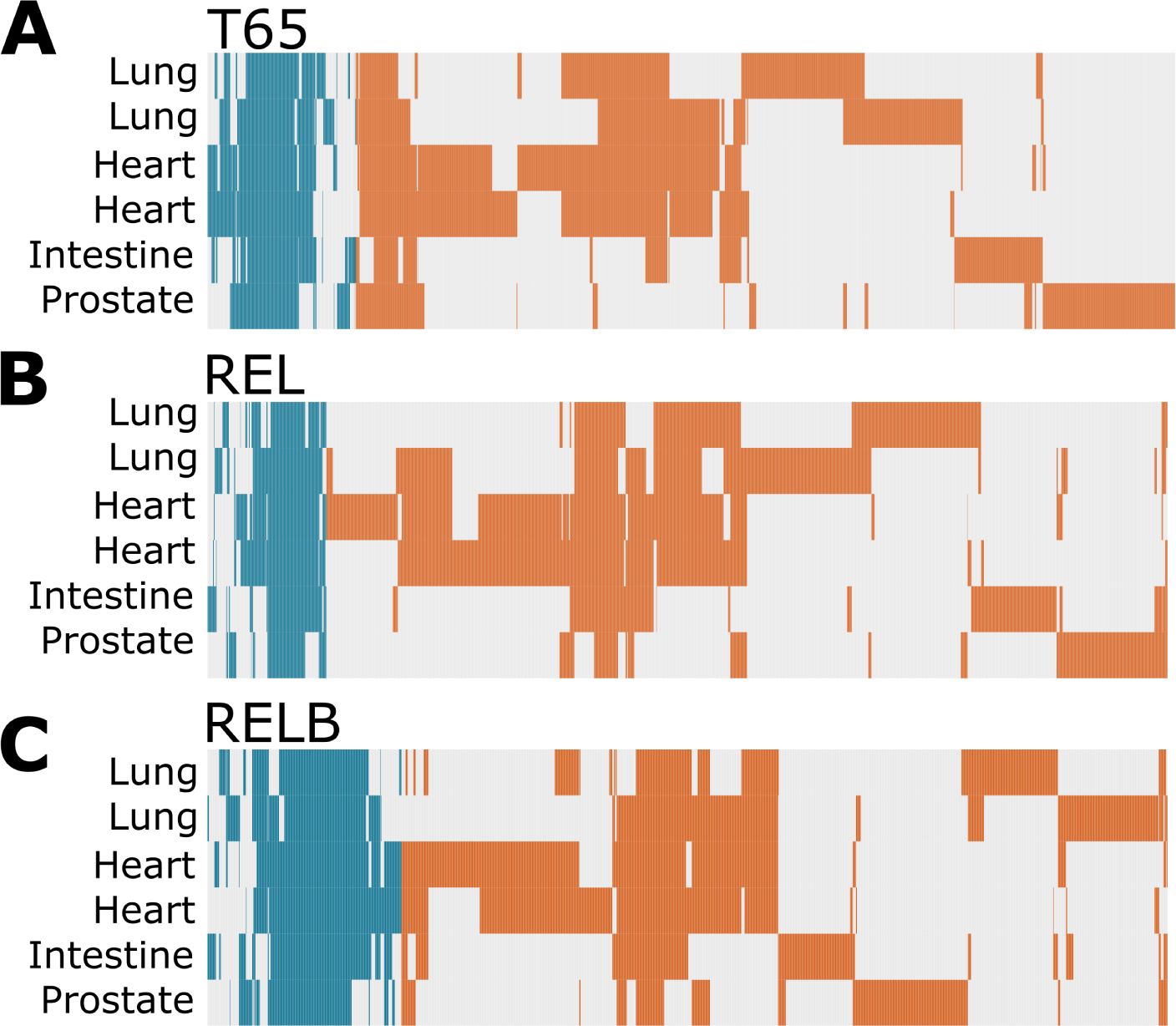
Additional NF*κ*B subunits. Similar to Figure 5C. Additional NF*κ*B subunit shown: A) T65, B) REL or C) RELB. Orange represents enhancer regions, the teal represents promoter regions. The absence of color in a given tissue indicates that that region does produce a bidirectional in that tissue type. Across all subunits, the majority of regions are enhancers (T65: 85.5%, REL: 87.6%, RELB: 79.8%) that are not shared across the distinct tissue types.

**Supplementary Figure 15.**
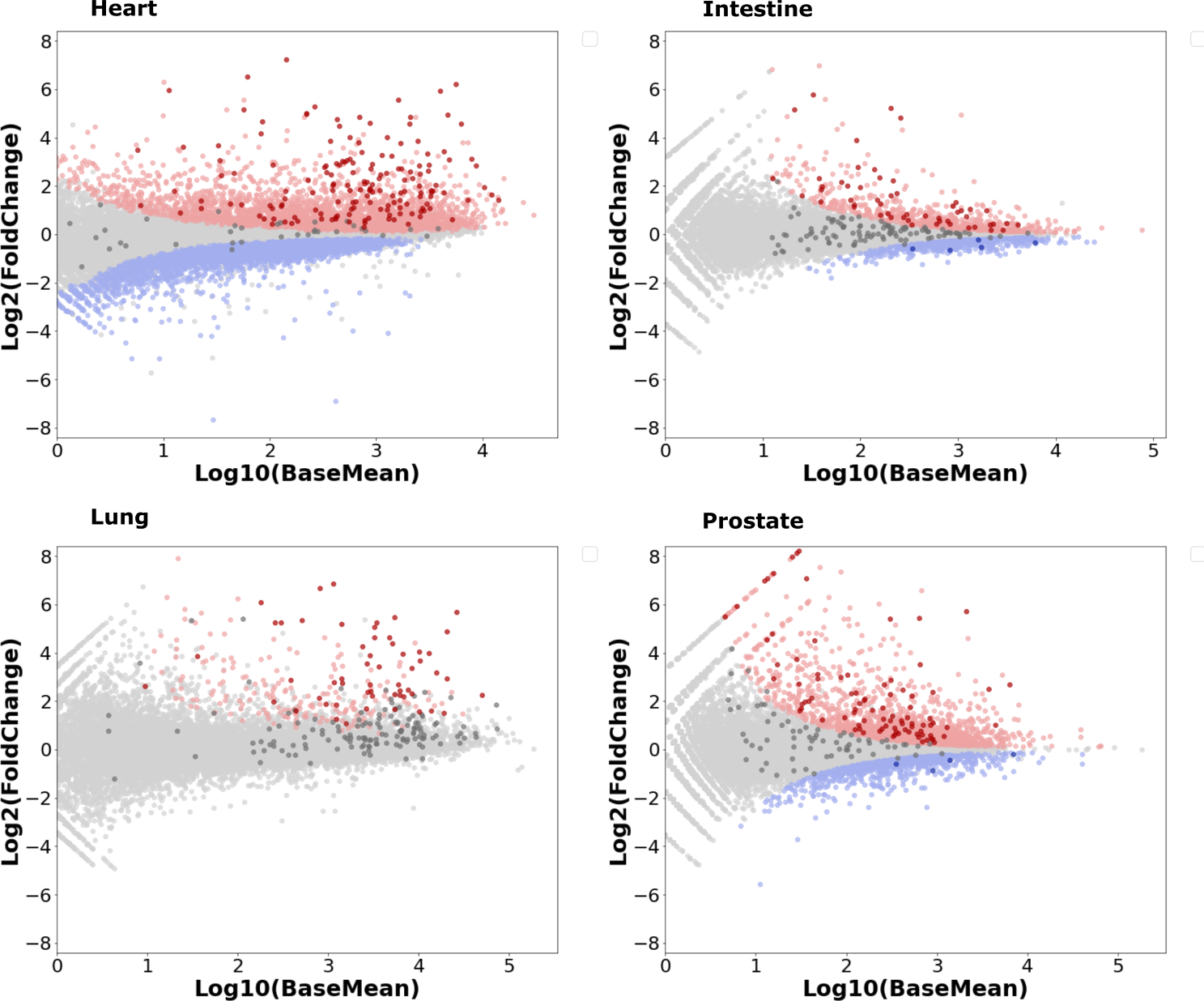
Overall NF*κ*B response is retained across tissue types. MA plots showing the differential expression results across the four tissues with TNF*α* treatment vs control[26, 66–70]. Red: significant up-regulated, blue: significant down-regulated (padj *≤*0.05) and in grey in non-significant genes. The darker colored points (red, blue and grey) are NF*κ*B target genes as defined by GSEA Hallmarks v5.0: TNF*α* signalling via NF*κ*B (n=200 genes). Note that many NF*κ*B target genes are significantly up-regulated across all tissues.

